# Cysteine Glutathionylation as a Global Dynamic Regulator of Protein Active Site Accessibility and Protein Complex Formation

**DOI:** 10.64898/2026.01.15.699703

**Authors:** Kish. R. Adoni, Nikolaos Tataridas-Pallas, Tianle Zhang, Fanindrah Deshmukh, Margaret Johncock, Oscar Shutt, Athanasios Bakalakos, Petros Syrris, Perry M. Elliott, John Labbadia, Konstantinos Thalassinos

## Abstract

Protein-glutathionylation is traditionally viewed as a protective mechanism that shields cysteine-residues from irreversible oxidative damage. Its broader functional roles remain poorly understood, in part due to technical limitations in detecting this modification at scale. Here, we develop and leverage a new mass-spectrometry approach that preserves protein-glutathionylation, thereby revealing its widespread distribution across the proteome in cell models, worms, mice and human cardiac tissues. In all cases, we find that glutathionylation sites are enriched at both protein-protein interfaces and protein-active sites, and are highly conserved across species. We find that glutathionylation is dynamically redistributed in response to environmental challenges, thereby driving remodelling of cellular protein–protein interaction (PPI) networks, and access to protein active sites, with functional and phenotypic consequences. Finally, we show that glutathionylation accumulates on key cardiac-sarcomeric proteins in aged-mice and human cardiomyopathy biopsies, revealing its role in cardiovascular dysfunction. These findings reposition glutathionylation as a crucial regulatory PTM, akin to phosphorylation, that orchestrates adaptive cellular responses. This work redefines the role of glutathionylation, with broad implications for cell biology and disease.

## Introduction

The fate of a cell is largely determined by the orchestrated spatiotemporal behaviour of its proteome in response to its environment. Protein Post-Translational Modifications (PTMs) can affect a protein’s structure and function, allowing rapid responses to cellular events^1^. Protein cysteine-PTMs represent an understudied biochemical modification, especially in a proteome-wide context. This is because mass spectrometry-based proteomics workflows, the preferred method for studying PTMs on a proteome-wide scale, include the reduction and alkylation of disulfide bonds as one of the first steps in sample preparation, which results in the loss of any cysteine PTMs that were present^2^. As such, the majority of available proteomics-PTM data in public data repositories^3^ underrepresent the occurrence of cysteine modifications.

There are, however, many modifications that can take place on cysteine residues^4^. The variable pKa of non-catalytic cysteine thiols can range between 7.4 and 9.1^5^. Subtle changes in the protein’s microenvironment can, therefore, trigger modifications in the oxidative status of the cysteine-thiol group. Cysteine S-glutathionylation, which forms from a disulfide bond between the protein and glutathione’s central cysteine R-group, is currently thought to protect cysteine thiols against oxidative damage from reactive oxygen species (ROS)^6,7^. Several studies have implicated protein glutathionylation in disease^8^, as well as in driving protein structural changes. Examples include the role of glutathione in modulating the conformation of titin^9^, Fatty Acid Binding Protein 5 (FABP5)^10^, Apoptosis-associated speck-like protein containing a CARD (ASC)^11^ and the gating of α-rings between open and closed forms of the 20S proteasome complex^12^.

Previous work in cysteine-modification based proteomics analyses has involved capping reduced cysteine residues followed by the incorporation of a reducing agent (e.g. dithiothreitol, DTT/ tris(2-carboxyethyl)phosphine, TCEP) and a second round of capping to reveal ratios of reduced/oxidised cysteines^13–16^. Further work has built on this concept with the use of advanced reduction/labelling strategies that contain enrichable handles for affinity purification^17^. Whilst effective in localising and quantifying cysteine oxidation, information pertaining to the specific PTM is lost with such strategies. More recently, biotinylated glutathione-based techniques have been used to affinity purify and identify glutathionylation sites^18^. As with many PTM enrichment strategies, these approaches introduce biases on account of adjacent amino acid interaction with the antibody^19–21^.

Here we describe the development of a new enrichment-free proteomics methodology for global cysteine-PTM identification, which has enabled us to identify and quantify over two-thousand glutathionylation sites across human bone osteosarcoma cells (U2OS), the nematode worm *Caenorhabditis elegans,* and mouse- and human-cardiac tissue (including biopsies from cardiomyopathy patients). These data revealed glutathionylation to be widespread across the proteome and to be dynamically regulated in response to changes in protein homeostasis, reduced insulin/insulin-like signalling and ageing. We show that cysteine residues modified by glutathionylation occur at sites that are involved in protein-protein interaction (PPI) interfaces, as well as in close proximity to protein active sites, and that such cysteines are highly conserved across species. Using whole-cell crosslinking mass spectrometry, we show that rewiring of PPIs correlates with changes in protein glutathionylation state. This was particularly evident across core components of the proteostasis network, influencing the ability of cells to counteract age-related protein aggregation and proteotoxicity. In addition, the level of active site glutathionylation was robustly altered across proteins from aged and diseased cardiac tissues.

Our work shows that glutathionylation is not just a means to protect proteins against oxidative stress but rather a dynamic and reversible modification, more akin to phosphorylation, that is capable of rewiring the organisation and functionality of the proteome in response to fluctuating physiological and environmental conditions.

## Results

### Protein glutathionylation is widely identified across the proteome

We have taken advantage of advancements in LC sensitivity, MS instrumentation and data processing workflows^22,23^, as well as developments in High Field Asymmetric Waveform Ion Mobility Spectrometry (FAIMS)^24^, to develop an enrichment- and label-free methodology for the identification of glutathionylation on a global scale (Figure 1A)

**Figure 1.**
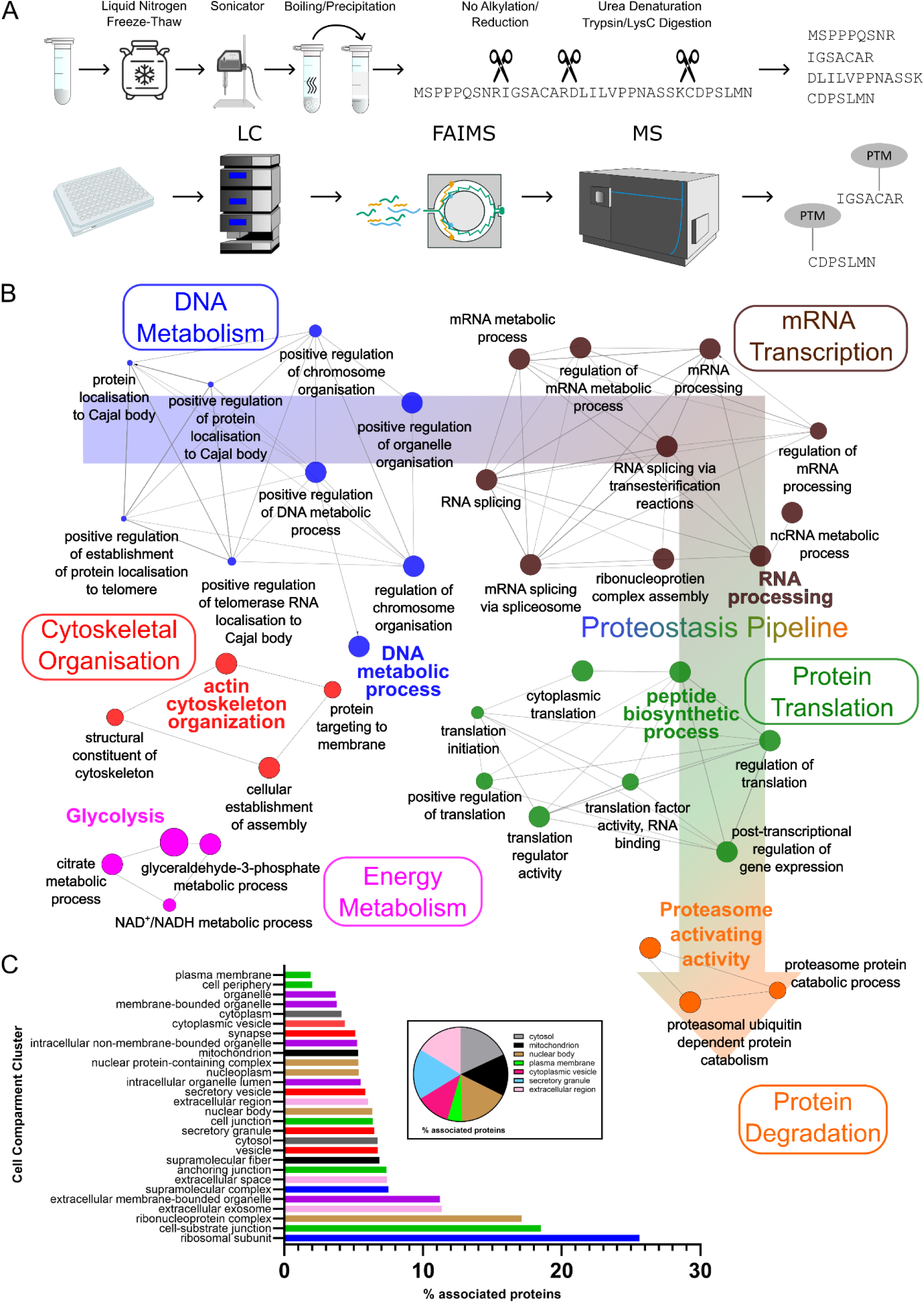
Protein glutathionylation is widely identified across the proteome. (A) Methodological workflow for glutathionylation-proteomics. (B) Cellular pathways associated with proteins identified in their glutathionylated proteoform. Circle sizes correspond to the number of proteins identified. (C) Cell compartment ontology analysis (GO:CC) revealed glutathionylated proteins to be found within the cytoplasm, nucleus and mitochondria, as well as secretory compartments including granules, vesicles and extracellular regions.

We initially subjected human osteosarcoma (U2OS) cells to our workflow and identified 22, 8, 13 and 16 cysteine -palmitoyl, -nitrosyl, -cysteinyl and -carbamyl PTMs respectively, as well as 898 cysteine glutathionylation sites corresponding to 565 proteins. This reveals protein-glutathionylation to be amongst the most common forms of cysteine-PTM found within cells (Figure S1A and B). Gene ontology analysis of glutathionylated proteins from U2OS cells revealed protein glutathionylation to be found in multiple cellular compartments and present on key factors involved in DNA metabolism (41 proteins), RNA processing (32 proteins), cytoskeletal organization (71 proteins), protein synthesis (76 proteins), protein folding (29 proteins) and protein degradation (13 proteins) (Figure 1B and C). In addition, and as expected based on previous reports^25^, 37 proteins related to energy metabolism (including glycolysis and aerobic respiration) were also found to be glutathionylated (Figure 1B). These data demonstrate that glutathionylation is widespread and occurs on proteins with critical roles in cell function and viability.

### Protein-glutathionylation preferentially occurs at protein-protein interfaces and protein active sites

Next, we decided to probe the structural and regulatory role of protein-glutathionylation in more detail. Using predicted protein structures from the AlphaFoldDB^26^, we found that glutathionylation sites are present on both buried and solvent exposed cysteines, with buried sites preferentially occurring near known protein active sites, and exposed sites being localised to solvent accessible protein surfaces (Figure 2A).

**Figure 2.**
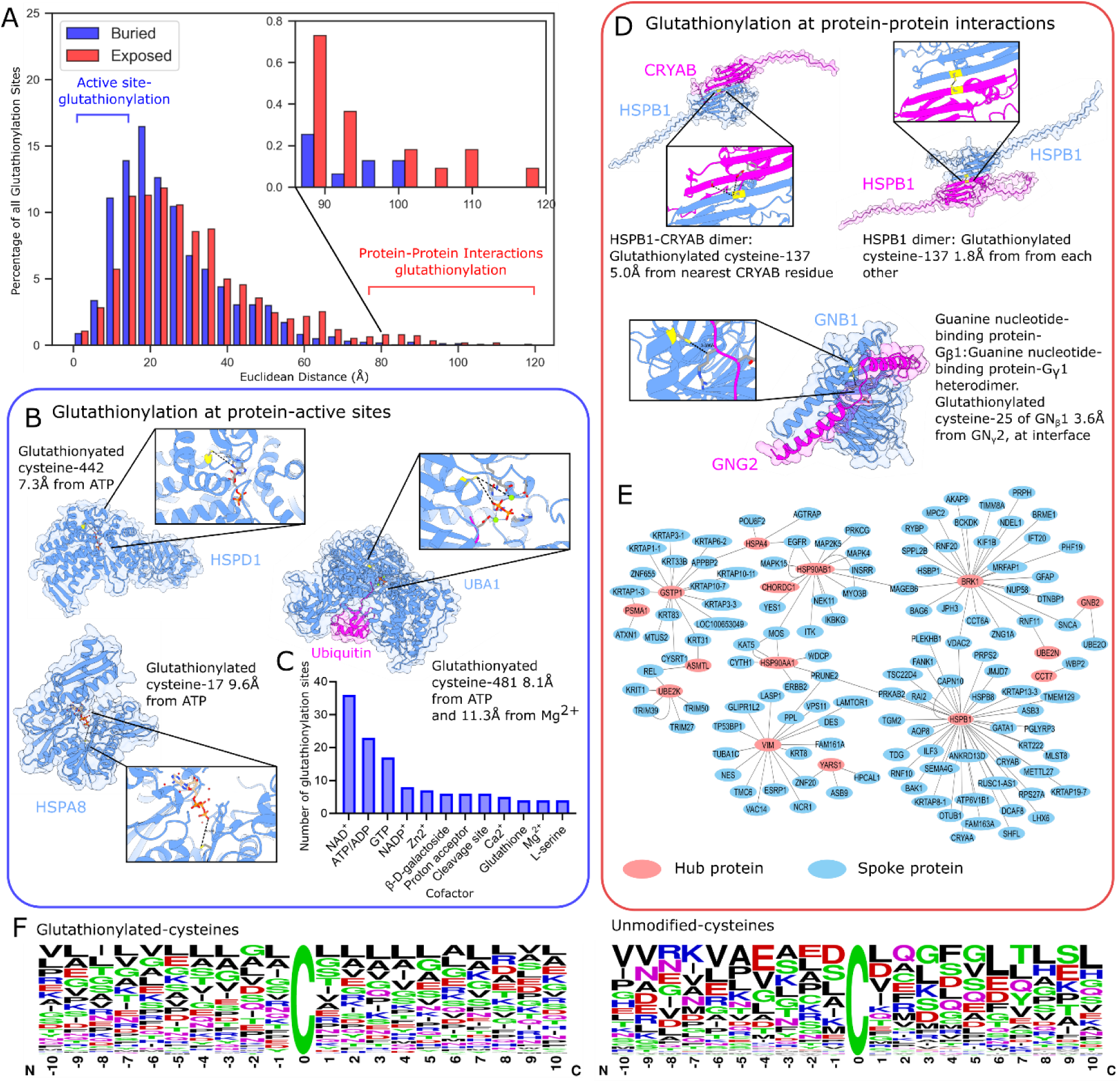
Protein-glutathionylation preferentially occurs at protein-protein interfaces and protein active sites. (A) Bar chart showing the Euclidean distance of protein-glutathionylation sites, relative to their nearest active-site. Glutathionylated residues were binned by their distance (0–120 Å, 10 Å bins) from the nearest catalytic site, revealing a bimodal pattern in which modifications cluster either adjacent to buried active sites or at solvent-exposed protein–protein interfaces. (B) Examples of glutathionylation sites adjacent to active sites, including: Heat Shock Protein-D1 (glutathionylated cysteine-442 7.3 Å from ATP at ATP active site) (PDBe: 8g7o^28^), Ubiquitin-like Modifier-Activating Enzyme-1 (glutathionylated cysteine-481 8.1 Å from ATP at ATP active site) (PDB: 6DC6^29^), Heat Shock Protein-A8 (glutathionylated cysteine-17 9.6 Å from ATP at ATP active site) (PDB: 6B1N^30^). (C) Bar chart corresponding to the number of glutathionylated cysteines adjacent (<10 Å from known active site) to a known active site. (D) Examples of glutathionylation sites at protein-protein interfaces, including: Heat Shock Protein-B1 heterodimeric interaction with Alpha-crystallin B chain (glutathionylated cysteine-137 at dimerization interface, 5.0 Å from nearest CRYAB residue), Heat Shock-Protein-B1 homodimer (glutathionylated cysteine-137 of both monomers at dimerization interface, 1.8 Å from each other) and Guanine nucleotide-binding protein-G_β_1 heterodimeric interaction with Guanine nucleotide-binding protein-G_γ_1 (glutathionylated cysteine-25 of GN_β_-1 3.6 Å from interface with GN_γ_-2). All dimeric structures generated from AlphaFold3^26^. (E) PPI network for all experimentally identified glutathionylation sites within 5 Å of a known heterodimeric interface. Proteins that were experimentally identified to be glutathionylated are annotated in red (generated with XlinkCyNET^31^ application within Cytoscape^32^). (F) Consensus motif analysis (WebLogo^33^) of glutathionylated cysteines vs non-modified cysteines.

Consensus motif analysis of the glutathionylated cysteine sites suggested that, whilst non-modified cysteines tended to be flanked by a combination of polar (serine, threonine), positively charged (arginine, lysine), negatively charged (glutamic acid) and hydrophobic (alanine, glycine, leucine and valine) residues, glutathionylated cysteine sites were preferentially flanked by more hydrophobic residues such as leucine, valine and alanine (Figure 2F). A comparison of the Euclidean distance of glutathionylated vs non-modified cysteines, suggests that glutathionylation occurs closer to protein active sites (Figure 2A), with 182 glutathionylated cysteines across 59 proteins found to be within 10 Å (Euclidean distance) of active sites^27^ associated with Nicotinamide Adenine Dinucleotide (NAD^+^), ATP/ADP, GTP/GDP, metal ion (Zn^2+^, Ca^2+^ and Mg^2+^), β-D-galactoside, glutathione, L-serine binding and protein cleavage (Figure 2C). Among these, we observed glutathionylation at cysteine residues close to the ATP binding sites of key protein quality control factors within the mitochondria (HSPD1/HSP60), cytosol and nucleus (HSPA8/HSC70 and UBA1) (Figure 2B).

### Protein Glutathionylation is a dynamic PTM that responds to environmental **stress**

To determine whether similar patterns of glutathionylation were observed *in vivo,* we applied our approach to the nematode worm *C. elegans*. Consistent with our findings in U2OS cells, glutathionylation occurred across the proteome and was observed across the same cellular machinery identified in U2OS cells (Figure 3A).

**Figure 3.**
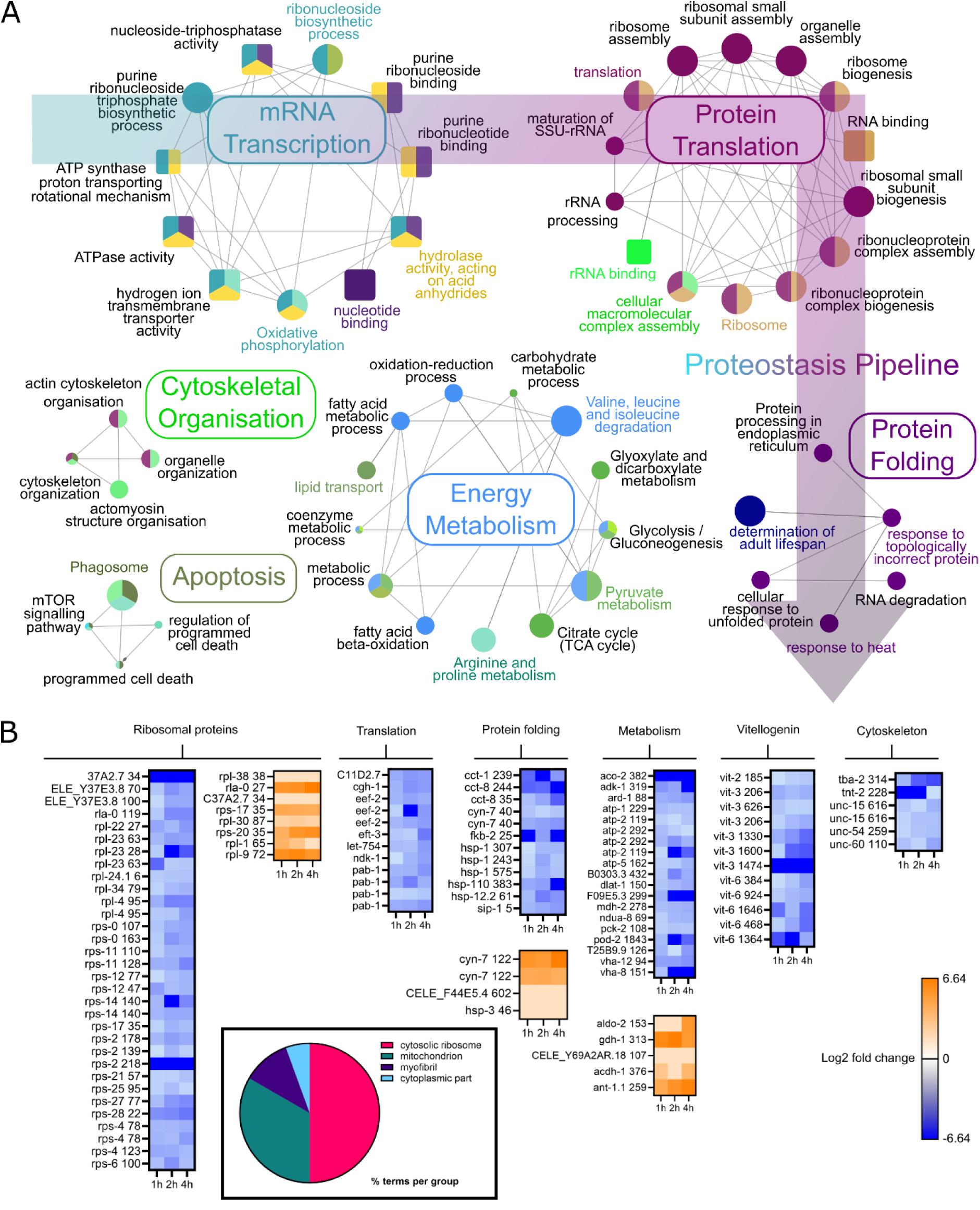
Protein Glutathionylation is a dynamic PTM that responds to environmental stress. (A) Biological processes and molecular functions associated with changes in protein glutathionylation following heat shock in *C. elegans* (day 1 adults; 35 °C). Squares represent molecular functions and circles represent biological processes. (B) Heatmaps corresponding to changes in the levels of glutathionylation (relative to non-heat shocked controls) across *C. elegans* proteins at 1-, 2- and 4-hours post heat-shock (35 °C). Pie chart denotes the cellular compartments that the glutathionylatable proteins are associated with.

We next asked whether glutathionylation patterns were dynamic and responsive to environmental challenges. To this end, we exposed worms to heat-shock (35 °C) for 1, 2 or 4 hours, and measured the levels of glutathionylation across the proteome. Our analysis revealed dynamic responses of glutathionylation sites to elevated temperature, with 75, 104 and 120 cysteine glutathionylation sites exhibiting increased or decreased abundance (log2 foldchange <-1.0 or >1.0) after 1, 2 and 4 hours of heat shock (Figure 3B). The majority of glutathionylation was decreased in response to heat stress (Figure S2B) and these changes were not simply due to the decreased abundance of proteins (Figure S2A). Together, our results show that protein-glutathionylation is dynamically remodelled across the proteome in response to environmental changes.

### Remodelling of protein-glutathionylation impacts protein homeostasis and **tissue health**

To complement our thermal stress experiments, we next investigated the impact of pharmacologically or genetically remodelling glutathionylation on cell and tissue health. To do this, we took advantage of buthionine sulfoximine (BSO), an inhibitor of γ-glutamyl cysteine synthetase mediated synthesis of reduced-glutathione^34^ and insulin/insulin-like receptor (DAF-2) defective mutants, which are known to exhibit differential expression of multiple glutathione-related genes, enhanced stress resistance and increased lifespan^35–38^.

Both BSO treatment and reduced insulin/insulin-like signalling resulted in the redistribution of glutathionylation sites across the proteome (Figure 4A and B, Figure S3A and B). Interestingly, the distribution of protein-glutathionylation following BSO treatment or mutation of *daf-2* was similar, with 50 and 194 glutathionylation sites identified in BSO and control conditions, respectively, and 19 and 102 glutathionylation sites increased/decreased in *daf-2* mutants, respectively (log2 FC >1.0 or <-1.0) (Figure 4A and B). Among the proteins exhibiting reduced glutathionylation in response to BSO treatment or mutation of *daf-2*, were multiple subunits of the T-complex Protein Ring Complex (TRiC), which is formed of two octameric rings stacked on top of each other that act to fold approximately 10% of the proteome in mammalian cells^39^. We identified 11 glutathionylation sites, distributed across seven of the eight TRiC subunits (CCT1, 2, 4, 5, 6, 7, and 8) (Figure S4A and B). Based on the human TRiC complex crystal structure (PDBe:7lup^40^), these glutathionylation events are predicted to interfere with interactions between CCT1/CCT4 (CCT1 cysteine-239 and -400), CCT2/CCT4 (CCT4 cysteine-410), CCT5/CCT7 (CCT5 cysteine-252 and -406), CCT7/CCT8 (CCT7 cysteine-29 and -509) and CCT8/CCT7 (CCT8 cysteine-35), thereby suppressing formation of the TRiC complex (Figure S4C). Interestingly, CCT7 cysteine-29, CCT8 cysteine-35 and CCT8 cysteine-244 exhibited reduced glutathionylation in response to BSO treatment, loss of DAF-2 activity and heat shock, suggesting that under these conditions, removal of glutathione is used to promote TRiC complex assembly and activity (Figure S4B).

**Figure 4.**
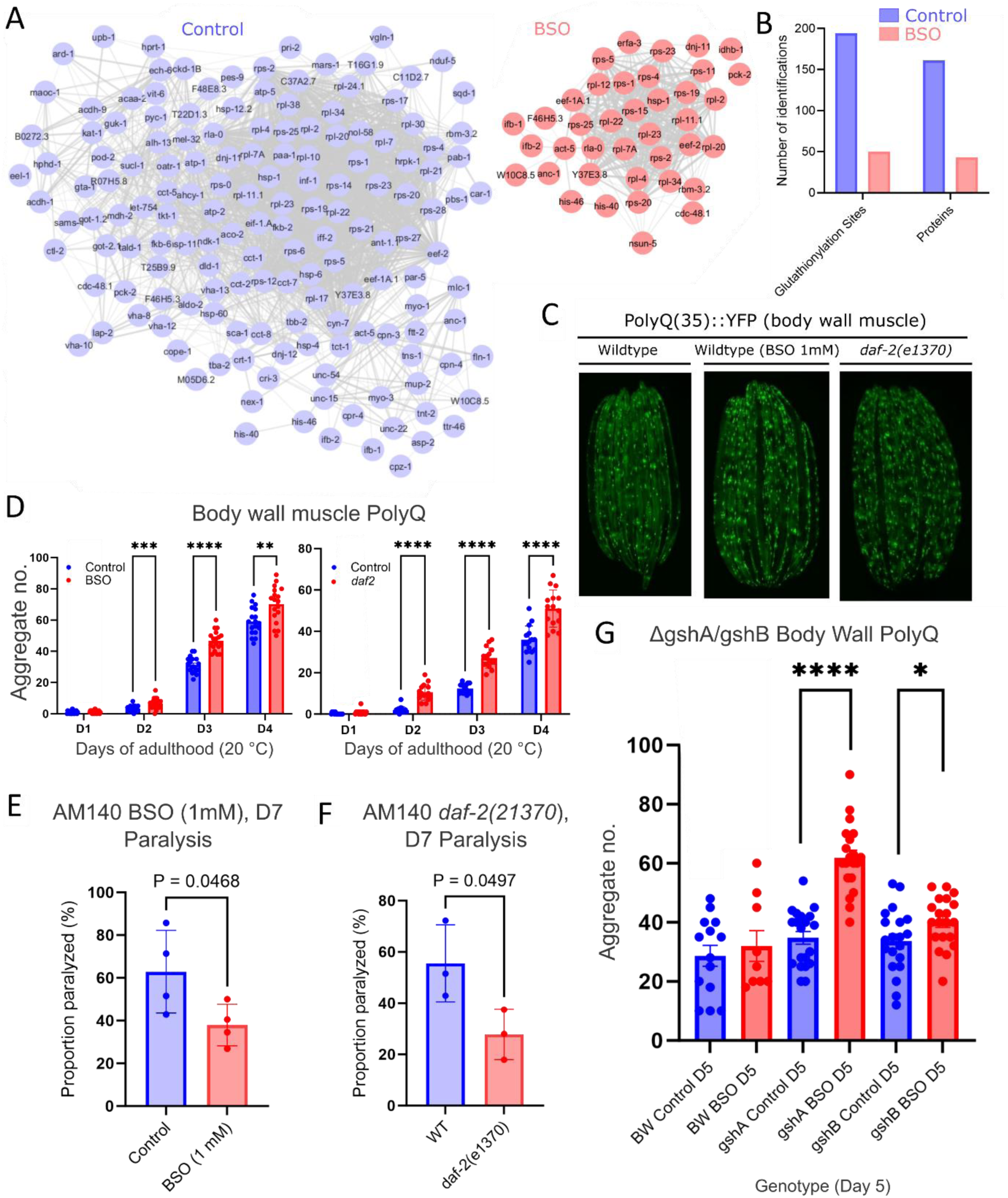
Remodelling of protein-glutathionylation impacts protein homeostasis and tissue health. (A) The glutathionylated protein network in control (blue) and BSO (1 mM) treated (red) *C. elegans*. All identifications were present in at least 2 of 3 biological replicates of the condition. (B) Bar chart illustrating the relative occurrence of protein-glutathionylation in control (blue bars) and BSO treated worms (red bars), in the context of glutathionylated proteins and glutathionylation sites. (C) Representative fluorescence microscopy images of polyQ aggregation in the body wall muscles of wild type (control and BSO) and *daf-2* mutant animals on day 3 of adulthood. (D) Numbers of protein aggregates in the body wall muscles of control (blue bars) and BSO treated (red bars) wild type and *daf-2* mutant worms, on different days of adulthood. (E, F) Relative levels of paralysis observed on day 7 of adulthood in (E) control or BSO treated worms, and (F) wild type and *daf-2* mutant worms, expressing polyQ(35)::YFP in body wall muscles. (G) Levels of polyQ(35)::YFP aggregation on day 5 of adulthood in the body wall muscles of worms grown on wildtype (BW) or glutathione synthesis defective (*gshA* and *gshB*) *E. coli* and treated with control or BSO (1 mM) conditions. Bars represent means +/- SD and statistical tests were performed using two-way ANOVA (d and g) or paired, two-tailed Student’s t-test (e and f). * = p < 0.05, ** = p < 0.01, *** = p < 0.001 and **** = p < 0.0001.

In total, 11 glutathionylated cysteine residues were identified across components of the TRiC complex, 8 of which map to CCT protein–protein interfaces. These interface cysteines exhibit strong evolutionary conservation: 5 are strictly conserved as cysteine in all organisms analysed (yeast to human), with the remaining 3 varying in only a single species. Conversely, cysteine residues outside the interfaces show weaker conservation across species. This pattern indicates that interface-associated cysteines are under stronger selective pressure than non-interface positions, and that glutathionylation/deglutathionylation events are a highly conserved mechanism for fine tuning TRiC protein complex assembly and proteome integrity. (Figure S4D).

Having consistently demonstrated the presence of glutathione on several key factors related to protein quality control, and the dynamic response of protein-glutathionylation in response to protein folding stress (i.e. heat shock), we next considered the functional implications of glutathionylation on protein homeostasis.

Polyglutamine fused to YFP (PolyQ::YFP) exhibits age-dependent protein aggregation and leads to cellular toxicity when expressed in *C. elegans* tissues^41–43^, thereby acting as a well-established sensor of proteostasis capacity. Both BSO treatment and reduced insulin/insulin-like signalling resulted in an increase in polyQ::YFP aggregate number in body wall muscle tissues (Figure 4C and D). Similar effects were also observed within intestinal cells following BSO treatment (Figure S3C and D) and these effects were enhanced when dietary ingestion of glutathione was prevented by growing worms on *E. coli gcsA* and *gcsB* mutants, which are unable to synthesize glutathione (Figure 4G). Surprisingly, the increased protein aggregation caused by BSO treatment or mutation of *daf-2*, was associated with a reduction in muscle paralysis (i.e. polyQ toxicity) (Figure 4E and F), suggesting that pharmacological or genetic remodelling of glutathionylation across the proteome promotes a protective form of protein aggregation that protects long-term tissue health.

### Glutathionylation influences tissue health by re-wiring protein-protein interactions

To understand how the remodelling of glutathionylation across the proteome suppresses polyQ toxicity in muscle tissues, we first looked for any proteins exhibiting changes in active site glutathionylation in both BSO treated and *daf-2* mutant worms. We identified 4 metabolic enzymes that consistently exhibited reduced glutathionylation following BSO treatment (T22D1.3/IMPDH, SAMS-1, GSPD-1 and GTA-1), with GSPD-1 (6-phosphogluconate dehydrogenase) also reduced in *daf-2* mutants and in response to heat shock (Figure 5A-C).

**Figure 5.**
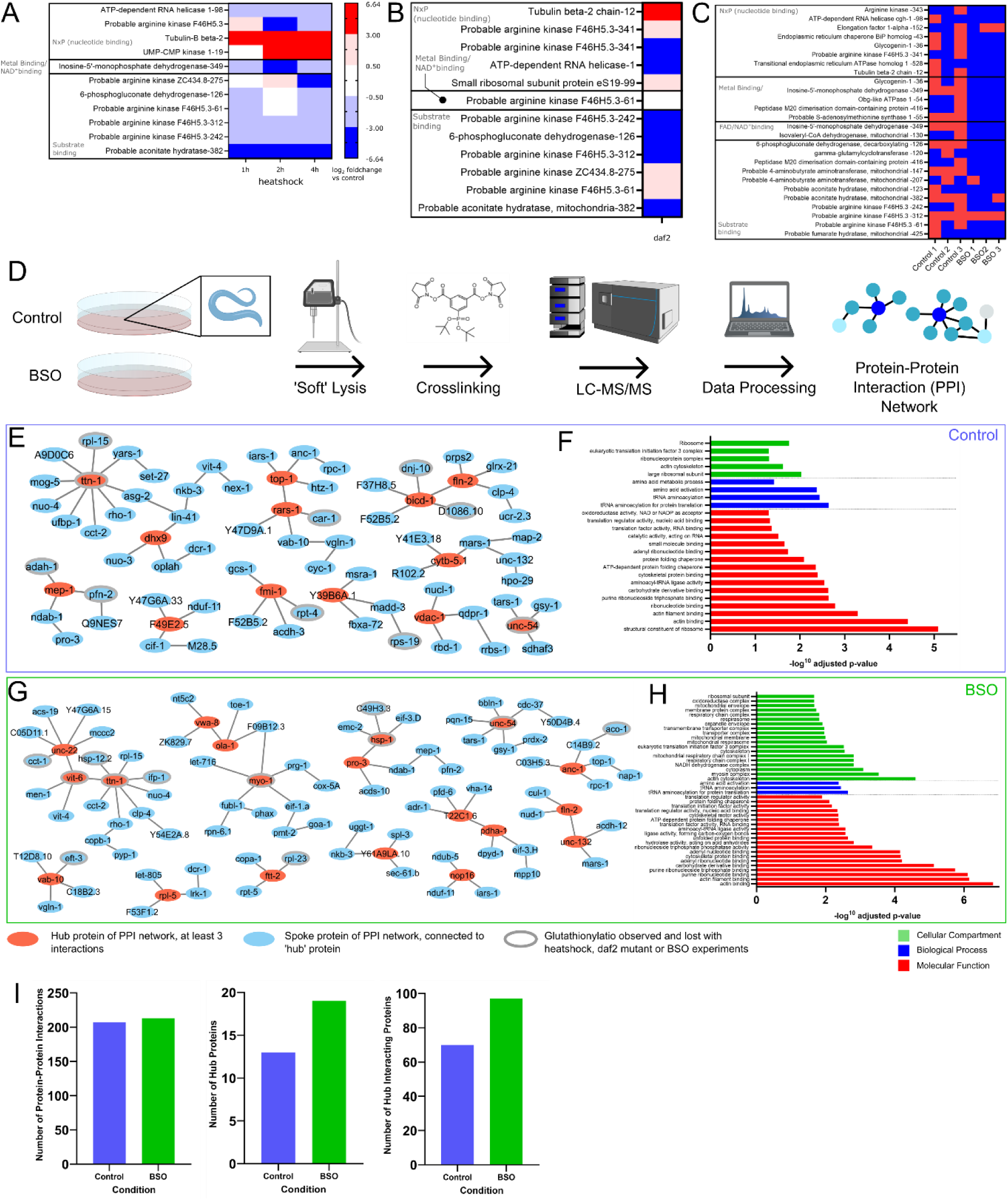
Glutathionylation influences tissue health by re-wiring protein-protein interactions. (A-C) Experimentally identified glutathionylation sites adjacent to known active sites (≤10 Å) from (A) heat shock treatment, (B) genetic perturbation (daf2 mutant) and (C) pharmacological treatment (BSO) experiments. (D) Workflow of *C. elegans* ‘soft’ lysis to maintain non-covalent PPIs before crosslinking and LC-MS/MS. (E-H) All ‘hub’ proteins (with three or more inter-protein interactions, red) and ‘spoke’ proteins (blue), and the associated gene ontology biological processes (blue), molecular functions (red) and cell compartments (green) for the proteins within the network for (E and F) control and (G and H) BSO treated *C. elegans*. Where a PPI was observed upon loss of glutathionylation (by BSO treatment, heat-shock and/or *daf2* mutant), the glutathionylated protein is bordered in grey. (I) The number of PPIs, number of hub proteins and number of ‘hub interacting (spoke) proteins for control and BSO treatment.

Given that glutathionylation also occurs within close proximity (5 Å) of predicted heterodimerisation or homodimerisation interfaces, we next considered whether the effects of BSO treatment on protein aggregation stem from glutathionylation-dependent rewiring of the cellular protein-protein interaction (PPI) network. Whole-cell lysate crosslinking mass spectrometry (Figure 5D) revealed that a similar number of PPIs were recovered across control (206) and BSO (212) conditions (Figure 5E-I). BSO treatment, however, resulted in a dramatic rewiring of PPIs, with a 46% increase in hub proteins undergoing three or more heteromeric interactions, and a 39% increase in hub interacting (spoke) proteins (Figure 5E-I).

By mapping protein glutathionylation onto the hub/spoke PPI networks, we found 11 PPIs that occurred exclusively in response to BSO treatment, and 15 hub/spoke PPIs that were disrupted by BSO treatment. Among these, HSP-1/HSC70 was identified as a hub protein exclusively in BSO treated animals (Figure 5E-H). Four HSP-1/HSC70 cysteine glutathionylation sites were downregulated by heat-shock, BSO treatment and/or *daf-2* mutation. These included cysteines at the interaction sites of HSP-1/HSC70 and C49H3.3/LLPH, Eukaryotic Elongation Factor-3 (EIF3), and ER membrane protein complex subunit 2 (EMC2) (Figure S5A-C). Of these glutathionylation sites, HSP-1 cysteine-575 was conserved from worms to humans, while HSP-1 cysteine-17 was conserved from *E. coli* to humans, further supporting the role of glutathionylation events as an ancient mechanism for modulating protein complex assembly dynamics (Figure S5D). ersytert

In addition to HSP-1/HSC70, we also observed many PPIs with titin-1, a large muscle protein predicted to associate with many proteins. Of the titin-PPIs identified, heat shock protein-12.2 (HSP-12.2), Intermediate filament protein-1, CLPF-1 and vitellogenin-6 were found to exclusively interact with titin-1 following BSO-treatment (Figure 5E-H). Furthermore, HSP12.2 cysteine-61 glutathionylation was reduced by heat-shock, BSO treatment and *daf-2* mutation, with BSO treatment driving the titin1-HSP-12.2 interaction in all three biological replicates (titin-1 and HSP-12.2 were not found to interact under control conditions, where HSP12.2 was found to be glutathionylated).

Finally, we also observed that reduced protein glutathionylation upon BSO treatment stimulated PPIs in the mitochondria (corresponding to respiratory complexes such as NADH dehydrogenase, respiratory chain I, mitochondrial respirasome and membrane protein complexes) and cytoplasm (ribosomal complex, eukaryotic translation initiation complex, and myosin/actin cytoskeletal complexes), whereas a gain in protein glutathionylation was associated with the formation of ribosomal, cytoskeletal and eukaryotic translation initiation complexes (Figure 5E-H). Overall, our analysis reveals that the protein-protein interactome is dramatically rewired in response to the redistribution of glutathionylation, consistent with our identification of steric hindrance of glutathionylation sites at protein-protein interfaces.

To determine whether glutathionylation-dependent changes in PPIs have functional relevance, we knocked down all proteins that were found to form PPIs in response to BSO treatment, concomitant with cysteine-site loss of glutathionylation, in our heat-shock, *daf-2* mutant and BSO treatment data sets (Figure 6A and B). While HSP-1/HSC70 exhibited changes in glutathionylation status in BSO treated worms and was identified as a key hub protein in our BSO induced PPI network, it was excluded from our candidate RNAi screen as it is already known to greatly increase polyQ aggregation and toxicity and cause anatomical and reproductive defects in *C. elegans* when knocked down^44–46^.

**Figure 6.**
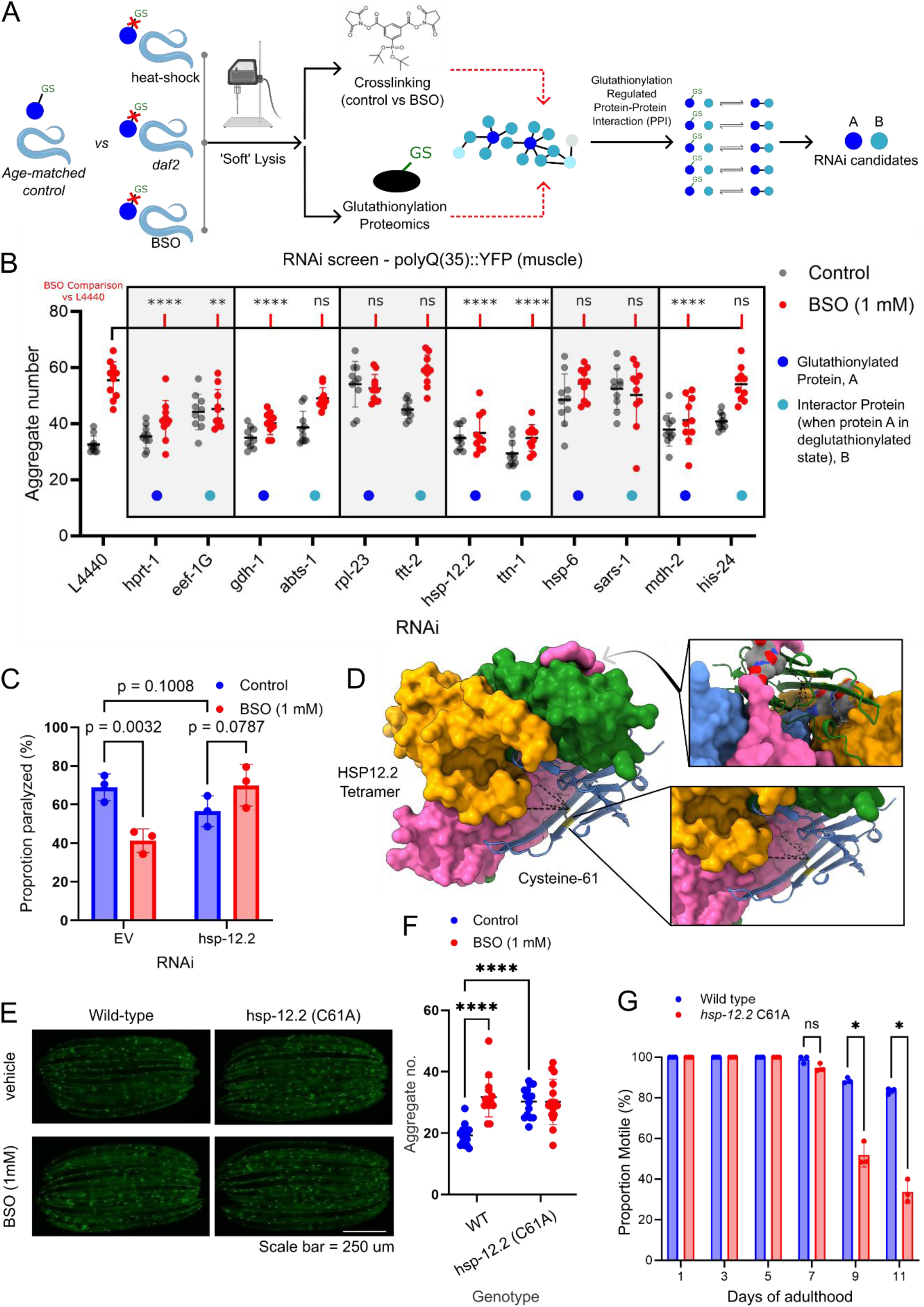
Glutathionylation of HSP12.2 regulates protein aggregation. (A) Filtering strategy for RNAi screen candidates. (B) Levels of polyQ(35)::YFP aggregates in body wall muscles on day 4 of adulthood following RNAi knockdown of candidate protein pairs in control or BSO treated worms. Thick line boxes represent proteins involved in the PPI. Dark blue circle represents glutathionylatable protein, and teal blue circle represents interacting protein. (C) Proportion of paralyzed worms on day 7 of adulthood in worms expressing polyQ(35)::YFP in body wall muscles and subjected to *hsp-12.2* RNAi under control or BSO conditions. (D) AlphaFold 3 generated HSP12.2 homodimer, with cysteine-61 <12 Å from 5 residues of its interacting subunit. (E) Representative fluorescence microscopy images of polyQ aggregation in the body wall muscles of wild type (control and BSO) and *hsp-12.2(C61A)* mutant animals on day 3 of adulthood. (F) Numbers of polyQ(35)::YFP aggregates in control and BSO treated wildtype and *hsp-12.2*(C61A) mutant worms. (G) Barchart to represent percentage of motile animals for control and *hsp-12.2(C61A)* mutant, from day 1 to day 11. Bars represent mean values +/- SD and statistical comparisons were performed by unpaired Student’s t-test (b) or two-way ANOVA (C, F and G).

RNAi against our remaining candidate protein pairs revealed that knockdown of several proteins influenced aggregate formation under control or BSO treatment conditions. Only in the case of the HSP12.2 - titin interaction pair, however, did knockdown of both interaction partners prevent BSO induced polyQ aggregation without increasing aggregation under control conditions as well (Figure 6B). Consistent with this, we found that cysteine-61 glutathionylation of HSP-12.2 is predicted to sterically hinder homotetramer formation (Figure 6C), client interaction and complex formation which are important for HSP-12.2 activity^47^. Mutation of cysteine 61 to alanine resulted in increased polyQ aggregation, comparable to BSO treatment in wildtype animals (Figure 6E and F). The constitutive absence of glutathionylation of HSP-12.2, accelerated muscle paralysis in wild type animals (Figure 6G). Together, these data suggest that protein-glutathionylation induced changes in PPIs have functional relevance to cell and tissue health. The fact that constitutively preventing HSP-12.2 glutathionylation accelerates age-related muscle decline, supports the notion that glutathionylation events are used to dynamically shift protein function, as and when required, to support cell and tissue health.

### Dynamic protein glutathionylation drives the redistribution of protein-protein interaction networks in mouse and human cardiac tissue

Having observed a functional relationship between glutathionylation-mediated protein-protein interactions (PPI) and cellular homeostasis, we next explored its role in mammalian tissue health. Given the large number of glutathionylated muscle-related proteins in our *C. elegans* dataset, as well as previous protein specific studies implicating glutathionylation in the function of human muscle proteins (e.g. glyceraldehyde phosphate dehydrogenase^48^, actin^49,50^, mitochondrial complex II^51^, ras^52^ sarco(endo)plasmic reticulum Ca2+ ATPase^53^, myosin binding protein C^54^) we investigated age-related dynamic protein-glutathionylation in cardiac tissue from mice at different life stages. Consistent with our U2OS and *C. elegans* data, an inverse relationship between protein glutathionylation and the extent of PPIs was evident. Cardiac tissue from 2-month-old mice exhibited the lowest levels of glutathionylation and a corresponding relative increase in proteins participating in the PPI network compared with older age groups (Figure S6).

We next considered whether these findings are relevant to the health of human cardiac tissue. To this end, we analysed myocardial tissue from a ‘control’ individual and myocardium resected during septal myectomy in 6 patients suffering from hypertrophic cardiomyopathy (6 with heterozygous pathogenic variants in *mybpc3* (3 patients), or *myh7* (1 patient) and Y gene elusive (2 patients) (Figure 7A). Principal Component Analysis of the global and glutathionylated proteome revealed that our control sample was distinct from any of the cardiomyopathy samples (Figure 7B). As expected, variation within the cardiomyopathy groups was observed with greatest similarities between *mybpc3* and gene elusive tissues (Figure 7B). By grouping the various cardiomyopathy patient-derived cardiac biopsies and comparing them to the control sample, protein glutathionylation was increased by 42%, with concomitant 29% decrease in the number of PPI network proteins (Figure 7C). Further, investigation of the PPI networks for the two conditions revealed a decrease in hub proteins from 18 (control) to 13 (cardiomyopathy). The decreased propensity for protein interactions was demonstrated by the reduced interaction network of tropomyosin-1, from 6 (control) to 2 (cardiomyopathy) PPI’s (Figure 7D-F). Further, tropomyosin-1 interaction with tropomyosin-3 was only observed in the control condition, with its glutathionylation at cysteine-190 located at the interface of the dimer (Figure 7F). Mapping of identified glutathionylation sites onto the cardiac sarcomeric thick and thin filament crystal structures revealed multiple examples of modifications at interfaces. As well as tropomyosin cysteine-190’s potential involvement in homo-dimerization, myosin-7 cysteine-1748 was glutathionylated at its interface with titin and myosin light chain-3 (MLC3) cysteine-182 was glutathionylated at its interface with myosin (Figure 7F). In addition, we found glutathionylation at cysteine-12 of the GTP coordinating site of beta-tubulin (also conserved in *C. elegans*), a crucial regulator of tubulin polymerization (Figure 7G).

**Figure 7.**
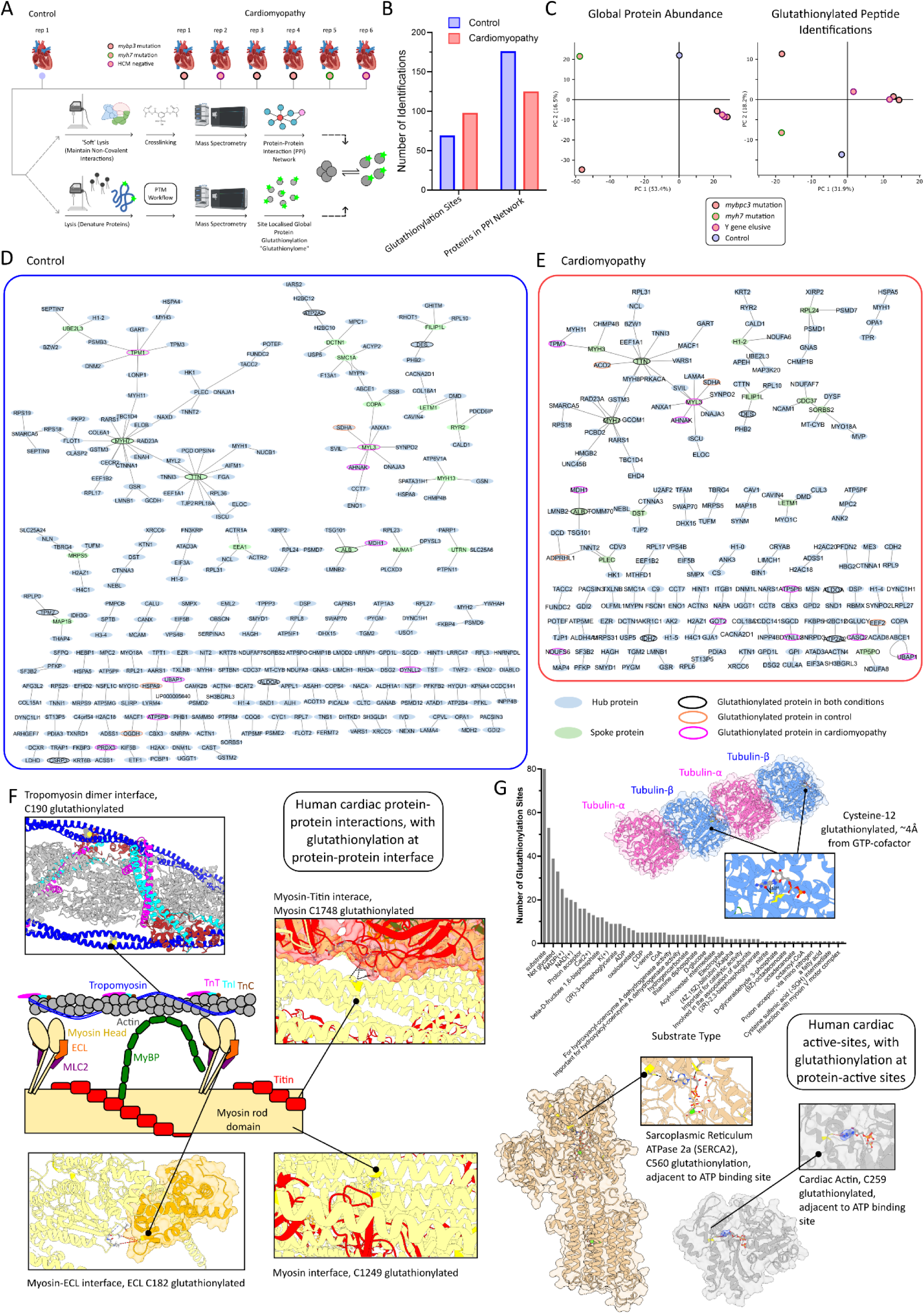
Dynamic protein glutathionylation drives the redistribution of protein-protein interaction networks in human cardiac tissue. (A) Workflow for glutathionylation proteomics and crosslinking MS of human cardiomyopathy patient biopsies. (B) Glutathionylation was upregulated in cardiomyopathy, compared to control samples, with concomitant reduction of proteins involved in PPI networks. (C) Principal Component Analysis for global proteomics and glutathionylome of six cardiomyopathy patient biopsies, including three patients with *mybpc* variants, two patients presenting with no cardiomyopathy-related variants (Y gene elusive), one patient presenting with *myh7* variant, and one control patient. All variants were heterozygous. (D, E) PPI network of control vs cardiomyopathy cardiac tissue from patient derived cardiac biopsies, annotated with hub proteins (green), spoke proteins (blue), proteins that were glutathionylated in control and cardiomyopathy presenting patients (black outline), proteins that were glutathionylated exclusively in control (orange outline) and proteins that were glutathionylated exclusively in cardiomyopathy presenting patients (pink outline). (F) Schematic representation of thick and thin filament of cardiac sarcomere, with glutathionylation sites of tropomyosin and actin mapped onto the crystal structure for the thin filament (pdbe: 6kn8^55^) and glutathionylation sites of myosin-7, myosin light chain-3 and titin mapped onto the crystal structure of thick filament (pdbe: 8g4l^56^). PPI interface with glutathionylation observed at tropomyosin dimer (cysteine-190), myosin-7 dimer (cysteine-1249), myosin-MLC3 (MLC3 cysteine 182) and myosin-titin (myosin-7 cysteine-1748). (G) Bar chart to show number of glutathionylation sites within 10Å of cofactor coordinating site, with crystal structures for co-localisation of glutathionylation with ATP-coordination site of cardiac actin and sarcoplasmic endoplasmic reticulum calcium ATPase 2 (SERCA2) (pdbe: 7bt2^57^) and GTP-coordination site of tubulin-beta.

Our initial observations in U2OS cells showed that many glutathionylation sites are not only located close to PPI interfaces but are also proximal to protein active sites (Figure 2A). Within cardiac tissue, co-localisation of glutathionylation with active sites (≤10Å Euclidean distance) was identified across 40 proteins and was associated with diverse co-factors, the most frequent being [2Fe-2S], electron carrier (e.g. NAD, FAD) and metal ion binding (Figure 7G). One notable example included glutathionylation of cysteine-259 at the ATP co-ordinating site of actin, which was exclusively observed within cardiomyopathy samples, and not in the control (Figure 7G).

## Discussion

As well as vastly increasing the functional repertoire of a protein, post-translational modifications are known to modulate the cellular protein-interaction network^58^, augmenting the cell with an additional, real-time spatio-temporal layer of global PPI regulation. Whilst protein-cysteine residues have long been associated with dynamic oxidative PTMs, LC-MS/MS, the gold standard for PTM identification, generally uses reduction/alkylation workflows that are not amenable to cysteine-PTM characterisation. In this work, we developed a workflow for the unbiased, unlabelled global identification of cysteine-PTM’s to identify and localise the widespread occurrence of cellular protein-glutathionylation.

Current consensus predicates protection of protein cysteine sulfhydryl residues against oxidative damage as the major functionality of protein glutathionylation^6^. Our findings transform our understanding of protein glutathionylation, demonstrating that it acts as a dynamic regulator of non-covalent protein-protein interactions (PPI) to rewire the cellular proteome network. The global scale of protein glutathionylation, and its dynamic response to changing environments, suggest this PTM to be regulated in a manner more akin to phosphorylation.

Bioinformatics analysis of our experimentally identified protein glutathionylation sites, within human osteosarcoma cells, revealed their preferential occurrence at cysteine residues that are adjacent to protein active sites, as has been reported for the protection of active site cysteines against oxidative damage^59,60^. The localisation of a large sub-population of glutathionylated cysteines to known protein-protein interfaces, the majority of which are highly conserved across species, suggested a regulatory role for this PTM in the rewiring of the cellular protein-protein interactome. Further, the dynamic response of these PPI interface-localised glutathionylation events, to proteostatic and metabolic stresses, were indicative of an orchestrated redistribution of the cellular PPI network, as it adapts to its environmental challenges. Our findings add credence to the notion that protein behaviour should be considered in the context of their combinatorial assemblies that contribute to protein interaction networks, rather than simply as isolated modules that act independently of each other^61^.

Of the 512 known heteromeric interactions that were predicted to be sterically regulated by our experimentally localised glutathionylation events, cellular proteostasis functionality was heavily implicated, including the TRiC complex, HSC70/HSPA8 and small heat shock protein interactions. Glutathionylation is likely to disrupt complex formation, client interaction and and/or access to ATP. As evidence of this, we discovered that preventing glutathionylation of HSP-12.2 enhances its ability to promote polyglutamine aggregation and bind to the muscle protein titin. sHSPs engage with and stabilize misfolded proteins, thereby sequestering them into aggregate structures that can later be disassembled and subjected to refolding by disaggregase and HSP70 complexes^62^. Our findings suggest that glutathionylation of sHSPs and HSP70 acts as an additional layer of regulation that could allow cells to rapidly shift the equilibrium between protein folding or holding strategies. Such a mechanism has been described for the phosphorylation of small heat shock proteins^63–67^, therefore, future work to understand the interplay between phosphorylation and glutathionylation in the regulation of sHSP activity will be important. Our findings raise the intriguing possibility that glutathionylation allows cells to direct/pause/prime chaperone complex assembly and functionality as and when required ^62,68,69^.

We found that glutathionylation also occurs on proteins that are important for muscle function in worms, mice and humans. Previous work has implicated protein glutathionylation of glyceraldehyde phosphate dehydrogenase^48^, actin^49,50^ and mitochondrial complex II^51^ in ischemia-reperfusion injury, and sarco(endo)plasmic reticulum Ca2+ ATPase (SERCA)(*60*) and ras in atherosclerosis and cardiac hypertrophy^52,70^. Furthermore, glutathionylation of myosin binding protein C (cMyBP-C) at cysteine-249 has been reported to attenuate its phosphorylation, with implications on Ca^2+^ activated force development in ventricular myocytes^54^. Our finding integrate these disparate findings into a complete picture of how changes in glutathionlylation affect heart health.

Co-factor protein binding is central to cell and tissue health with the hydrolysis of ATP essential for muscle function. Previous work has linked the level of glutathionylation of G-actin to its polymerisation rate in A431 cells^50^. Similarly, glutathionylation of actin at cysteine-374 has been proposed to abrogate its polymerisation^71^. Our findings provide a functional role for the steric hindrance of actin binding with ATP, thus shifting the soluble/filamentous actin equilibrium towards actin ‘catastrophe’ and providing a mechanism for previous observations relating to actin-glutathionylation. Further, cysteine-259 glutathionylation was exclusively observed within cardiomyopathy samples, and not in the control, suggesting the propensity for dysregulation of actin polymerisation in sarcomeric tissue of cardiomyopathy patients. The same mechanistic principles were observed for tubulin polymerisation, where we identified glutathionylation of cysteine-12 at the GTP co-ordinating site of tubulin-beta^72^. We also observed glutathionylation on the N (nucleotide) domain of SERCA2 in mouse and human heart tissue. SERCA2’s active transport of Ca^2+^ against its concentration gradient, into the sarcoplasmic reticulum to maintain the Ca^2+^ membrane gradient that controls cardiac excitation/contraction coupling, is dependent on its hydrolysis of ATP. Thus, our findings suggest a link between SERCA2 glutathionylation and cardiac contraction. To this end, SERCA2 glutathionylation has previously been associated with changes in intracellular Ca^2+^ concentration, with its dysregulation being associated with atherosclerosis^53,73^.

Our work has focused on pathways related to protein homeostasis and muscle function. Our data, however, reveal that glutathionylation is prevalent across multiple cellular pathways with remarkable conservation from cells to worms to mammals and is strongly implicated in cardiac tissue health. Given that our proof of principle experiments in worms reveal that a single glutathionylation event can dramatically alter cellular function and tissue health, future work to understand its role in other diseases, especially those that results from protein homeostasis collapse is important.

As with phosphorylation, an extensive network of enzymes^74^ (glutathione transferases and glutaredoxins ^75^ dictates the levels of glutathionylation across the cell. Future work to comprehensively describe how these enzymes respond to different stresses and define which clients they target will be key to manipulating the cellular ‘glutathionylome’ for therapeutic gain.

Ultimately, this work proposes protein glutathionylation as a major dynamic regulator for the redistribution of non-covalent protein-ligand and PPIs, to rewire the cellular proteome interaction network as the cell adapts to its environment (Figure S6).

## Resource Availability

### Supplementary content

- Supplementary Figures
- Dataset of all glutathionylation identifications in U2OS, *C.elegans* heat shock, daf2 mutant and BSO treatment experiments and crosslinking experiment.
- The mass spectrometry proteomics data have been deposited to the ProteomeXchange Consortium via the PRIDE^3^ partner repository with the dataset identifier PXD065262. Username, Password: FqKtzL33WXNg.

## Acknowledgements

The work was funded by BBSRC grants BB/T013273/1 and BB/P005535/1 to J.L and a Wellcome Trust Multiuser Equipment grant (221521/Z/20/Z) and Wellcome Trust Collaborative Award in Science (209250/Z/17/Z) to K.T. We thank Dr. Georgina Charlton for culturing U2OS cells, and Professor Gillian Bates (UCL IoN) for providing mouse cardiac tissues used in this work.

## Author Contributions

Conceptualization, K.R.A., J.L., K.T.

Methodology K.R.A.

Software T.Z., M.J, K.T.

Investigation K.R.A., N.T., T.Z., F.D. O.S.

Writing - Original Draft K.R.A.

Writing - Review & Editing K.R.A., J.L, K.T.

Visualization K.R.A

Supervision J.L., K.T.

Project administration K.T.

Funding acquisition J.L., K.T.

## Competing interests

*The authors declare no competing financial interest*.

## Methods

### C. elegans strains and maintenance

*C. elegans* were maintained at 20°C on polystyrene plates containing nematode growth medium (NGM) and OP50 *E. coli*^76^. Worms were routinely transferred to new plates under a Leica TL3000 stereo dissecting microscope using a platinum wire pick to avoid starvation or overcrowding. The following strains were used: wild type (N2; Bristol), AM140 (rmIs132[*unc-54*p::polyQ(35)::yfp]), AM738 (rmIs297[*vha-6*p::polyQ(44)::yfp]), JPL8 (*daf-2(e1370);*rmIs132*[unc-54p::polyQ(35)::yfp]*), PHX9765 (*hsp-12.2 (syb9765)*) and JPL86 (rmIs132[*unc-54*p::polyQ(35)::yfp];*hsp-12.2 (syb9765)*). The syb9765 mutant was generated by SUNY Biosciences and results in a cysteine to alanine conversion at residue 61 (C61A).

### Treatment with BSO and RNA interference

Buthionine sulfoximine (BSO) was dissolved in water to make a 100 mM working stock. This was added to the bacterial lawn of NGM OP50 plates to give a final concentration 1 mM BSO. Plates were then allowed to dry for 2 days at room temperature before being used for experiments. RNA interference was performed using bacterial clones obtained from the Vidal library^77^. RNAi bacteria were grown overnight (37°C, 180 rpm) in LB media containing 100 ug/ml ampicillin. Following this, cultures were induced for 3 hours (37°C, 180 rpm) with 5 mM IPTG and then seeded onto NGM plates containing 1 mM IPTG and 100 ug/ml ampicilin. Seeded RNAi plates were allowed to dry for 3 days at room temperature before being used for experiments.

### Imaging and quantifying polyglutamine aggregates

Worms were age-synchronized by egg-lay, grown to the desired stage of adulthood and then moved to unseeded NGM or RNAi plates for imaging. Worms were transferred into a drop of 0.5M sodium azide and arranged prior to imaging using an eyelash pick. Images were then captured using a Nikon SMZ1270 fluorescence dissecting microscope with a Nikon DS-Fi3 camera and NIS Elements L imaging software. To score aggregate load, worms were immobilized and imaged as above. Images were opened using FIJI/ImageJ and the number of puncta was recorded for each worm under different treatment conditions. At least 10 worms were scored per group, and counting was performed blind to treatment.

### Measuring polyglutamine toxicity in body wall muscles

Worm motility was used as a measure to assess polyglutamine toxicity in body wall muscles. Animals were age-synchronized by egg-lay and allowed to grow to adulthood. At the desired days of adulthood, worms were transferred into a small circle (5 mM diameter) drawn outside of the bacterial lawn on fresh NGM OP50, BSO or RNAi plates and allowed to move freely at 20°C for 1 hour. Worms that had not fully escaped the circle after 1 hour were touched with a platinum wire pick up to three times. Any worms that did not move forwards or backwards at least 1 body length when touched, were scored as paralysed.

### Worm lysis and protein extraction

Approximately 5000 worms were collected in M9 buffer^76^ and snap frozen in liquid nitrogen. Worms were then thawed on ice, re-suspended in RIPA buffer, and subjected to lysis using ceramic beads in a Precellys 24 homogenizer. Worms were checked under a stereo dissecting microscope to ensure they were fully destroyed (i.e., no intact tissues/carcasses were detected).

### Statistics and software

Bar charts to illustrate toxicity and aggregate number were generated in Prism v10.4.2^78^. Statistical test for toxicity was 2-way ANOVA, and for aggregate number was 2-way ANOVA with Turkeys method. Gene Ontology Enrichment Analysis was performed using STRING^79,80^, using default settings, and visualised within Cytoscape^32^. Molecular Function/Biological Process graphics were generated using the ClueGO^81^ app within Cytoscape, with default settings. XlinkCyNET^31^ app within Cytoscape was used for visualisation of PPI networks. All bar charts were generated within GraphPad Prism 10.4.2. For analysis of glutathionylated proteins in U2OS cells, the glutathionylation site was identified in both biological replicates (2/2) and at least one PSM corresponding to the glutathionylation site had a Byonic score >300. For analysis of glutathionylated proteins in *C. elegans* from the heat shock and *daf-2(e1370)* mutant experiments, at least one PSM corresponding to the glutathionylation site had a Byonic score >300. For analysis of proteins with dynamic glutathionylation that is lost upon treatment with BSO, and concomitantly gains a protein-protein interaction (PPI), the threshold was set such that the protein must be glutathionylated in at least 2 of 3 control biological replicates, and the crosslink was identified in at least two of three BSO-treated biological replicates. All *C. elegans* proteomics experiments were run in biological triplicate. For mouse cardiac samples, two biological replicates per condition were analysed, corresponding to 2-montth old, 12-month old and 20-month old. All reported glutathionylation sites were present in both biological replicates and all reported crosslinks were present in both biological replicates. For human biopsies of cardiac tissue, biopsies from six patients presenting with cardiomyopathy, including three *mybpc3* mutation carrying patients, two patients with no cardiomyopathy containing mutations (Y gene elusive) and one *myc3* mutation carrying patient, were analysed, as well as one control patient. All patients with reported mutations were heterozygous. Validated glutathionylation sites and PPIs were present in at least 2 of 6 cardiomyopathy patient biopsies.

Sequence alignment analysis was performed using blastp suite from the National Laboratory of Medicine, National Centre for Biotechnology Information (NCBI), with database: reference proteins (refseq proteins), algorithm: blastp (protein-protein blast)^82^.

### Bioinformatics

AlphaFold-predicted protein structures were retrieved from the AlphaFold Protein Structure Database (AlphaFold DB)^83–85^. We used the PDB module from Biopython library to query 404 human protein entries (by UniProt ID) against AlphaFold DB, obtaining 396 structures (8 IDs had no available model)^86^. Solvent-accessible surface area (SASA) for each glutathionylation site was calculated using FreeSASA, a C library with a Python module for SASA estimation^87^. Atomic radii were set as 1.6 Å for N, 1.7 Å for C, 1.4 Å for O, and 1.8 Å for S. Each modified cysteine was classified as buried if SASA ≤ 7 Å, or exposed if SASA > 7 Å. Functional site annotations (such as enzymatic active sites and ligand-binding residues) were obtained from the UniProt Knowledgebase for each protein^88^. These annotations provided the position of the functional sites used in subsequent analysis.

Distances between glutathionylated cysteine residues and nearby functional sites were computed in three-dimensional space. We used Biopython PDB module to calculate the Euclidean distance between the thiol S of each glutathionylated cysteine and the Cα atom of each annotated functional residue (For functional residue, we take thiol S instead of Cα atom)^86^. This allowed assessment of the spatial proximity of glutathionylation to known functional sites.

Known PPIs for the 396 human proteins were retrieved from Uniprot^88^. For each interacting protein pair, a predicted complex structure was obtained from the AlphaFold3 server^89,90^. Structural models of 4,609 binary complexes were analyzed using Biopython’s MMCIF parser and NeighborSearch module^86^. We measured the minimum distance between all atoms of each glutathionylated cysteine and all atoms in the interacting protein chain, and glutathionylation sites with the minimum distance ≤ 5 Å were considered at PPI interface.

Bioinformatics code is accessible on GitHub repository: https://github.com/TerryZhang175/PPI-tool.

### *Caenorhabditis elegans* MS sample preparation

For all *Caenorhabditis elegans* experiments, 3 biological replicates for each condition were analysed. Two liquid nitrogen freeze-thaw/homogenise cycles were performed with homogenising carried out using a 1.5 mL capacity plastic pestle before addition of an equal volume of lysis buffer (50mM Tris-Cl, 150mM NaCl, 50mM NaF, 25mM ß-glycerophosphate, 1% SDS, 1mM Ethylenediaminetetraacetic acid (EDTA), pH 7.6, 1x PhosSTOP phosphatase inhibitor per 50mL and 1x Pierce™ Protease Inhibitor Mini Tablets per 10mL), on ice. Samples were then probe sonicated (50W, 2 minutes, 10s on/off cycles) before supernatant extraction by centrifugation (16,000g, 10min). Freshly made urea (8M), 50mM ammonium bicarbonate and 1mM calcium chloride were added before incubation with shaking (4⁰C, 30min). Protein quantitation was performed using Rapid Gold BCA Protein Assay. 200µg of protein was taken for methanol-chloroform precipitation, by addition of 4 volumes of methanol, 1 volume of chloroform and 3 volumes of dH_2_O before vortexing and centrifugation (10,000g, 5 min, room temperature). Supernatant was discarded and 3 volumes of methanol was added before vortexing and a second centrifugation (10,000g, 5 min, room temperature) followed by removal of supernatant and air-drying of protein-pellet. Pellet was then reconstituted in digestion buffer (100mM Tris-Cl, 1% sodium deoxycholate (*w/v*), pH 8.0) followed by bath-sonication for 5 minutes. Trypsin/LysC was added at a protein:enzyme ratio of 25:1 (*w/w*) before incubation at 37⁰C with shaking for 4h, followed by 1:8 (*v/v*) dilution with 50mM Tris-Cl, pH 8.0 and overnight (16h) incubation. Digestion was quenched with trifluoroacetic acid (0.5 % *v/v*) before centrifugation (10,000g, 10 min) and extraction of supernatant followed by drying *in-vacuo*. Samples were then reconstituted in 4% acetonitrile, 0.5% trifluoroacetic acid before desalting with Oasis C18 HLB 96-well plate. Samples were then dried *in-vacuo* before reconstitution in 0.1% formic acid to a final concentration of 1mg/mL.

### *Caenorhabditis elegans* MS analysis

#### Heat shock and daf2 mutant experiments

An UltiMate 3000 RSLCnano liquid chromatography system (Thermo Fisher Scientific), with a 50 cm μPAC Neo HPLC analytical column (COL-NANO050NEOB) and a 0.075 × 20 mm trap cartridge (Acclaim PepMap C18 100 Å, 3 μm), was connected to a SilicaTip emitter. Column temperature was set at 45°C, and column flow rate was set to 750 nL/min. Mobile phase A (0.1% formic acid) and mobile phase B (100% acetonitrile, 0.1% formic acid) were applied with an elution gradient from 5.0 to 35.0% mobile phase B over 21 minutes. Total run time per sample was 30 minutes. FAIMS (High Field Asymmetric Waveform Ion Mobility Spectrometry) Pro was installed between the LC and MS, with internal FAIMS compensation voltages stepping across specific voltages: [−40V, −45V, −50V, −55V] and [−60V, −65V, −65V, −70V] over a 3 second cycle, as separate runs. FAIMS was set up in standard resolution with carrier gas flow rate set to 3.5 L/min. Analysis was performed using a Thermo Fisher Scientific Orbitrap Eclipse Tribrid mass spectrometer. The mass spectrometer was externally calibrated using a Pierce FlexMix calibration solution. nanoESI was performed by the application of a voltage to a SilicaTip emitter via an HPLC liquid junction cross. Spray stability and intensity was optimized by varying the SilicaTip electrospray voltage (1–2.5 kV) and varying SilicaTip positioning (in the *x*, *y*, and *z* dimensions). Transfer capillary temperature was set to 275 °C, RF lens was set to 40%, precursor ion mass spectra were acquired at a resolution of 60,000 with a mass range of 350 to 2000 *m*/*z* and a precursor ion charge state range of 2+ to 8+. MS1 spectra were recorded using data-dependent analysis with a cycle time of 3s across all four FAIMS CVs, automatic gain control was set to a target of 400,000, maximum injection time mode was set to Auto; precursor ions were isolated with a 1.6 *m*/*z* window using the quadrupole mass filter. For Orbitrap mass analysis based MS2 acquisition, higher-energy collision-induced dissociation was applied with a normalized energy of 30%. Orbitrap resolution was set to 30,000, scan range was set to Auto, Absolute AGC value was set to 50,000 and maximum injection time mode was set to Auto.

Data was processed using Proteome Discoverer 2.5, with the PMi Byonic node and searched against FASTA file *Caenorhabditis elegans* Reference proteome Proteome ID: UP000001940 downloaded from UniProt (September 2023). Precursor mass tolerance: 10ppm, Fragment mass tolerance: 20ppm, maximum missed cleavages: 3, Total common modifications max: 4, Total rare modifications: 1, Modifications: Oxidation (+15.994915Da, M, common 2), Cysteinyl (+119.004099Da, C, common 2), Glutathione (+305.068156Da, C, common 2), Carbamyl (+43.005814, C, common 2), Nitrosyl (+28.990164, C, common 2), Palmitoyl (+238.229666, C, common 2) and Acetyl (+42.010565, N-terminus, common 1), FDR: 0.01 (High confidence). Quantitation was performed using Label-Free Quantitation with Minora Feature Detector node and Feature Mapper node (all parameters set to default).

#### BSO treatment experiments

An UltiMate 3000 RSLCnano liquid chromatography system (Thermo Fisher Scientific), with a 50 cm μPAC Neo HPLC analytical column (COL-NANO050NEOB) and a 0.075 × 20 mm trap cartridge (Acclaim PepMap C18 100 Å, 3 μm), was connected to a SilicaTip emitter. Column temperature was set at 45°C, and column flow rate was set to 750 nL/min. Mobile phase A (0.1% formic acid) and mobile phase B (100% acetonitrile, 0.1% formic acid) were applied with an elution gradient from 5.0 to 35.0% mobile phase B over 48 minutes. Total run time per sample was 60 minutes. FAIMS Pro was installed between the LC and MS, and samples were run with external FAIMS fractionation at compensation voltages: −45V, −50V, −55V, −60V, −65V, −70V, −75V, −80V, −85V and −90V. FAIMS was set up in standard resolution with carrier gas flow rate set to 4.6 L/min. Analysis was performed using a Thermo Fisher Scientific Orbitrap Eclipse Tribrid mass spectrometer. The mass spectrometer was externally calibrated using a Pierce FlexMix calibration solution. nanoESI was performed by the application of a voltage to a SilicaTip emitter via an HPLC liquid junction cross. Spray stability and intensity was optimized by varying the SilicaTip electrospray voltage (1–2.5 kV) and varying SilicaTip positioning (in the *x*, *y*, and *z* dimensions). Transfer capillary temperature was set to 275 °C, RF lens was set to 40%, precursor ion mass spectrum was acquired at a resolution of 120,000 with a mass range of 350 to 2000 *m*/*z* and a precursor ion charge state range of 2+ to 8+. MS1 spectra were recorded using data-dependent analysis in Top20 mode, automatic gain control was set to a target of 400,000, maximum injection time mode was set to Auto, precursor ions were isolated with a 1.4 *m*/*z* window using the quadrupole mass filter. For Orbitrap mass analysis based MS2 acquisition, higher-energy collision-induced dissociation was applied with a normalized energy of 30%. Orbitrap resolution was set to 30,000, scan range was set to Auto, Absolute AGC value was set to 50,000 and maximum injection time mode was set to Auto.

Data was processed using Proteome Discoverer 2.5, with PMi Byonic node and searched against FASTA file *Caenorhabditis elegans* Reference proteome Proteome ID: UP000001940 2023_09_26.fasta downloaded from UniProt. Precursor mass tolerance: 10ppm, Fragment mass tolerance: 20ppm, maximum missed cleavages: 3, Total common modifications max: 4, Total rare modifications: 1, Modifications: Oxidation (+15.994915Da, M, common 2), Glutathione (+305.068156Da, C, common 2), Phospho (+79.966331, [S, T, Y], common 2), Acetyl +42.010565, [K, R], common 2) and Methyl (+14.01565, [K, R], common 2), FDR: 0.01. Quantitation was performed using Label-Free Quantitation with Minora Feature Detector node and Feature Mapper node (all parameters set to default).

#### Crosslinking MS sample preparation

Two liquid nitrogen freeze-thaw/homogenise cycles were performed with homogenisation carried out using a 1.5 mL capacity plastic pestle before addition of an equal volume of soft lysis buffer (20mM HEPES, 0.01% SDS (*v/v*), pH 7.6, 1x PhosSTOP phosphatase inhibitor per 50mL and 1x Pierce™ Protease Inhibitor Mini Tablets per 10mL) on ice. Sample was then probe sonicated (50W, 2 minutes, 10s on/off cycles) followed by supernatant extraction by centrifugation (16,000g, 10min). Protein quantitation was performed using Rapid Gold BCA Protein Assay. tert-Butyl Disuccinimidyl Phenyl Phosphonate (tBu-PhoX/TBDSPP) was equilibrated to room temperature for 15 minutes before making a 50mM stock solution in DMSO. TBDSPP was added to 1mg of sample (2mg/mL) to a final concentration of 1mM and samples were incubated at 4°C for 30 minutes, with the addition of 250uL of soft lysis buffer at 15 minutes. Crosslinking was quenched with ammonium bicarbonate to a final concentration of 20mM before the addition of 20uL alkaline phosphatase (37°C, 30min, shaking at 500rpm) to 200µg of sample. Sample was then reconstituted in 500µL of SDS lysis buffer (100mM Tris-Cl, 5% SDS, 5mM TCEP, 20mM chloroacetamide, pH 8.0) before incubation at 99°C for 10 minutes with shaking (1300rpm), cooling to room temperature and bath sonication for 10 minutes. Chloroform precipitation was performed as previously described to extract protein before the addition of digestion buffer (100 mM Tris-Cl, 1% sodium deoxycholate (*w/v*), pH 8.5) followed by vortexing and bath sonicating the protein pellet for 2 minutes to partially solubilise the precipitated protein. Trypsin (Promega Trypsin Gold) was added at a 1:50 enzyme:protein ratio and sample was incubated overnight (16h) at 37°C with shaking (1200rpm). Digestion was quenched with trifluoroacetic acid to a final concentration of 2% and incubated for 1h at 37°C with shaking (1200rpm) to deprotect the crosslinker. Samples were then reconstituted in 4% acetonitrile, 0.5% trifluoroacetic acid before desalting with Oasis C18 HLB 96-well plate. Samples were then dried *in-vacuo* before phosphoenrichment using Thermo Scientific High-Select TiO2 Phospho Enrichment Kit (Catalogue number: A32993) and following manufacturer instructions. Finally, sample was reconstituted in 0.1% formic acid at an approximated concentration of 1mg/mL for onward LC-MS/MS analysis.

#### Crosslinking MS analysis

An UltiMate 3000 RSLCnano liquid chromatography system (Thermo Fisher Scientific), with a 50 cm μPAC Neo HPLC analytical column (COL-NANO050NEOB) and a 0.075 × 20 mm trap cartridge (Acclaim PepMap C18 100 Å, 3 μm), was connected to a SilicaTip emitter. Column temperature was set at 45°C, and column flow rate was set to 300 nL/min. Mobile phase A (0.1% formic acid) and mobile phase B (100% acetonitrile, 0.1% formic acid) were applied with an elution gradient from 3 to 17.5% mobile phase B over 38 minutes then 17.5% to 35% over 10 minutes. Total run time per sample was 60 minutes. Analysis was performed using a Thermo Fisher Scientific Orbitrap Eclipse Tribrid mass spectrometer. The mass spectrometer was externally calibrated using a Pierce FlexMix calibration solution. nanoESI was performed by the application of a voltage to a SilicaTip emitter via an HPLC liquid junction cross. Spray stability and intensity was optimized by varying the SilicaTip electrospray voltage (1–2.5 kV) and varying SilicaTip positioning (in the *x*, *y*, and *z* dimensions). Transfer capillary temperature was set to 275 °C, RF lens was set to 40%, precursor ion mass spectrum was acquired at a resolution of 120,000 with a mass range of 350 to 2000 *m*/*z* and a precursor ion charge state range of 2+ to 8+. MS1 spectra were recorded with a data-dependent analysis in Top20 mode; automatic gain control was set to a target of 400,000; maximum injection time mode was set to Auto; precursor ions were isolated with a 1.4 *m*/*z* window using the quadrupole mass filter. For Orbitrap mass analysis based MS2 acquisition, higher-energy collision-induced dissociation was applied with a normalized energy of 30%. Orbitrap resolution was set to 30,000; scan range was set to Auto; Absolute AGC value was set to 50,000 and maximum injection time mode was set to Auto.

Data was processed using Proteome Discoverer 2.5, with Sequest and XlinkX nodes and searched against a bespoke FASTA file containing all identified proteins from the heat shock experiments. Precursor mass tolerance: 10ppm, fragment mass tolerance: 0.02Da, maximum missed cleavages was set to 2, Max equal dynamic modifications per peptide was set to 3, and max dynamic modification per peptide was set to 4, dynamic modifications included: Oxidation (+15.995Da, M), Acetylation (+42.011Da, N-terminus) and static modifications included Carbamidomethylation (+57.021Da, C). Default parameters were used for XlinkX nodes with the same FASTA file and modifications as for the Sequest node and the CSM FDR threshold and crosslink FDR threshold were set to 0.01. For a crosslink to be considered for onward analysis, it satisfied the criteria of being present in at least 2 of 3 biological replicates of at least one condition.

### U2OS

#### MS sample preparation

Two biological replicates were analysed. Cell pellet was harvested and lysed using 1:1 (*v/v*) ratio of lysis buffer (50mM Tris-Cl, 150mM NaCl, 50mM NaF, 25mM ß-glycerophosphate, 1% SDS, 1mM Ethylenediaminetetraacetic acid (EDTA), pH 7.6) with incubation at 4 °C for 30 minutes followed by centrifugation (16,000g) at 4 °C for 30 minutes. Supernatant was extracted and protein concentration was estimated by Bradford analysis^91^. Sample was then denatured by addition of freshly made 8M urea, co-administered with 50 mM ammonium bicarbonate and 1 mM calcium chloride. 50 mM Tris-HCl (pH 8.0) was then added before incubation of the sample at 37 °C with shaking, with Trypsin/LysC mix (purchased from Promega) at a 25:1 protein: enzyme ratio (*w/w*) for 4 hours followed by addition of 50 mM Tris-HCl (pH 8.0) to an eight-fold dilution. Sample was left to digest overnight (16h) before digestion was quenched with 0.5 % trifluoroacetic acid. Sample was then desalted using Pierce C18 Tips (purchased from Thermo Fischer Scientific) and reconstituted in 0.1 % formic acid at a concentration of 1 µg/µL before 5 µL of sample was injected for LCMS analysis.

#### MS analysis

An UltiMate 3000 RSLCnano liquid chromatography system (Thermo Fisher Scientific), with a 50 cm μPAC Neo HPLC analytical column (COL-NANO050NEOB) and a 0.075 × 20 mm trap cartridge (Acclaim PepMap C18 100 Å, 3 μm), was connected to a SilicaTip emitter. Column temperature was set at 45°C, and column flow rate was set to 300 nL/min. Mobile phase A (0.1% formic acid) and mobile phase B (100% acetonitrile, 0.1% formic acid) were applied with an elution gradient from 5.0 to 35.0% mobile phase B over 48 minutes. Total run time per sample was 60 minutes. FAIMS Pro was installed between the LC and MS, and samples were run with external FAIMS fractionation at compensation voltages: −30V, −35V, −40V, −45V, −50V, −55V, −60V, −65V, −70V, −75V, −80V, −85V and −90V (the data from these FAIMS CVs was then used to guide future LC-FAIMS-MS/MS glutathionylation experiments). FAIMS was set up in standard resolution with carrier gas flow rate set to 3.5 L/min. Analysis was performed using a Thermo Fisher Scientific Orbitrap Eclipse Tribrid mass spectrometer. The mass spectrometer was externally calibrated using a Pierce FlexMix calibration solution. nanoESI was performed by the application of a voltage to a SilicaTip emitter via an HPLC liquid junction cross. Spray stability and intensity was optimized by varying the SilicaTip electrospray voltage (1–2.5 kV) and varying SilicaTip positioning (in the *x*, *y*, and *z* dimensions). Transfer capillary temperature was set to 275 °C, RF lens was set to 40%, precursor ion mass spectrum was acquired at a resolution of 60,000 with a mass range of 350 to 2000 *m*/*z* and a precursor ion charge state range of 2+ to 8+. MS1 spectra were recorded with a data-dependent analysis in Top20 mode, automatic gain control was set to a target of 400,000, maximum injection time mode was set to Auto, precursor ions were isolated with a 1.6 *m*/*z* window using the quadrupole mass filter. For Orbitrap mass analysis based MS2 acquisition, higher-energy collision-induced dissociation was applied with a normalized energy of 30%. Orbitrap resolution was set to 30,000, scan range was set to Auto, Absolute AGC value was set to 50,000 and maximum injection time mode was set to Auto.

Data was processed using Proteome Discoverer 2.5, with PMi Byonic node and searched against FASTA file *Homo sapiens* Reference proteome downloaded from UniProt (July 2022). Precursor mass tolerance: 10ppm, Fragment mass tolerance: 20ppm, maximum missed cleavages: 3, Total common modifications max: 4, Total rare modifications: 1, Modifications: Oxidation (+15.994915Da, M, common 2), Cysteinyl (+119.004099Da, C, common 2), Glutathione (+305.068156Da, C, common 2), Carbamyl (+43.005814, C, common 2), Nitrosyl (+28.990164, C, common 2), Palmitoyl (+238.229666, C, common 2) and Phospho (+79.966331, [S, T, Y], common 2), FDR: 0.01 (high confidence). Quantitation was performed using Label-Free Quantitation with Minora Feature Detector node and Feature Mapper node (all parameters set to default). For a glutathionylation site to be considered for onward analysis, it was present in both biological replicates.

#### Mouse cardiac tissue and biopsies from patients presenting with cardiomyopathy

Sample preparation was performed as described in Section: *Caenorhabditis elegans* MS sample preparation with modification of the sample homogenisation process. After liquid-nitrogen freeze-thaw, a Dounce homogenizer, 1mL (Active Motif) was used to crush the cardiac tissue, with the addition of 250 uL of lysis buffer (20mM HEPES, 0.01% SDS (*v/v*), pH 7.6, 1x PhosSTOP phosphatase inhibitor per 50mL and 1x Pierce™ Protease Inhibitor Mini Tablets per 10mL), before proceeding with probe sonication as described in *Caenorhabditis elegans* MS sample preparation. Samples were then split into glutathionylation and crosslinking MS samples, with “*Caenorhabditis elegans* MS sample preparation” methodology followed for glutathionylation analysis, and “Crosslinking MS sample preparation” methodology followed for crosslinking MS. MS acquisition and data processing was performed as described for *C. elegans* glutathionylation proteomics and crosslinking MS, with the modification of FASTA file to *Mus musculus* (downloaded from UniProt on 25/08/2024) and *Homo sapiens* (downloaded from Uniprot on 20/07/2022). For mouse cardiac samples, two biological replicates per condition were analysed, corresponding to 2-montth old, 12-month old and 20-month old. All reported glutathionylation sites were present in both biological replicates and all reported crosslinks were present in both biological replicates. For human biopsies of cardiac tissue, biopsies from six patients presenting with cardiomyopathy, including three *mybpc3* mutation carrying patients, two patients with no cardiomyopathy containing mutations (HCM negative) and one *myc3* mutation carrying patient, were analysed, as well as one control patient. All patients with reported mutations were heterozygous. Validated glutathionylation sites and PPIs were present in at least 2 biological replicates of cardiomyopathy patient biopsies.

*For investigations on humans, informed consent was obtained from all participants after the nature and possible consequences of the studies were explained. The study was approved by the Health Research Authority Research Ethics Committee, ‘London - Bromley Research Ethics Committee’ under the title “Inherited Cardiac Disease: A Clinical and Genetic Investigation” (REC reference: 15/LO/0549; IRAS project ID: 167619; Amendment number: SA002; Amendment date: 14 October 2022)*

## Notes

### Competing Interest Statement

The authors have declared no competing interest.

