## Supplemental Figures for "Cysteine Glutathionylation as a Global Dynamic Regulator of Protein Active Site Accessibility and Protein Complex Formation"

### Supplemental Information

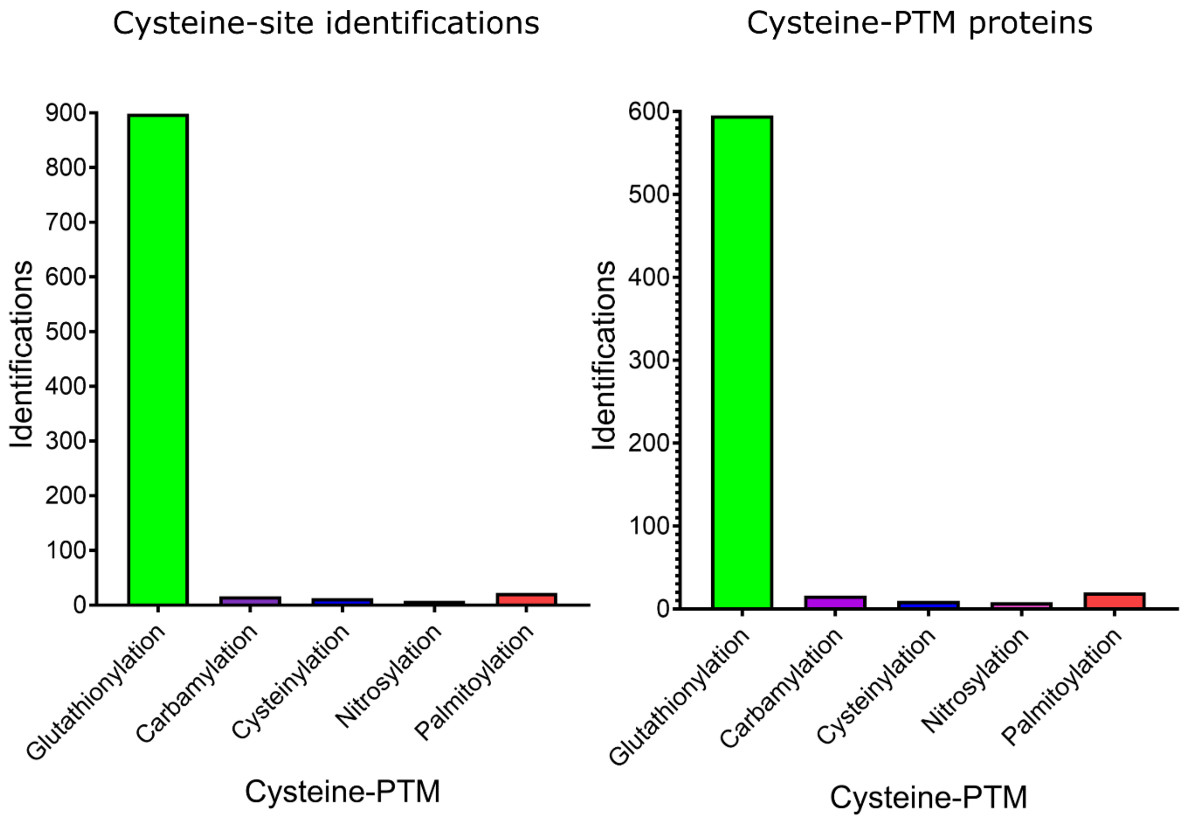


Figure S1. **Protein Glutathionylation is widely identified across the proteome.**

(A) Number of cysteine-PTM identifications including: glutathionylation, carbamylation, cysteinylation, nitrosylation and palmitoylation, using our cysteine-PTM proteomics workflow.

(B) Number of cysteine-PTM containing proteins including: glutathionylation, carbamylation, cysteinylation, nitrosylation and palmitoylation, using our cysteine-PTM proteomics workflow.

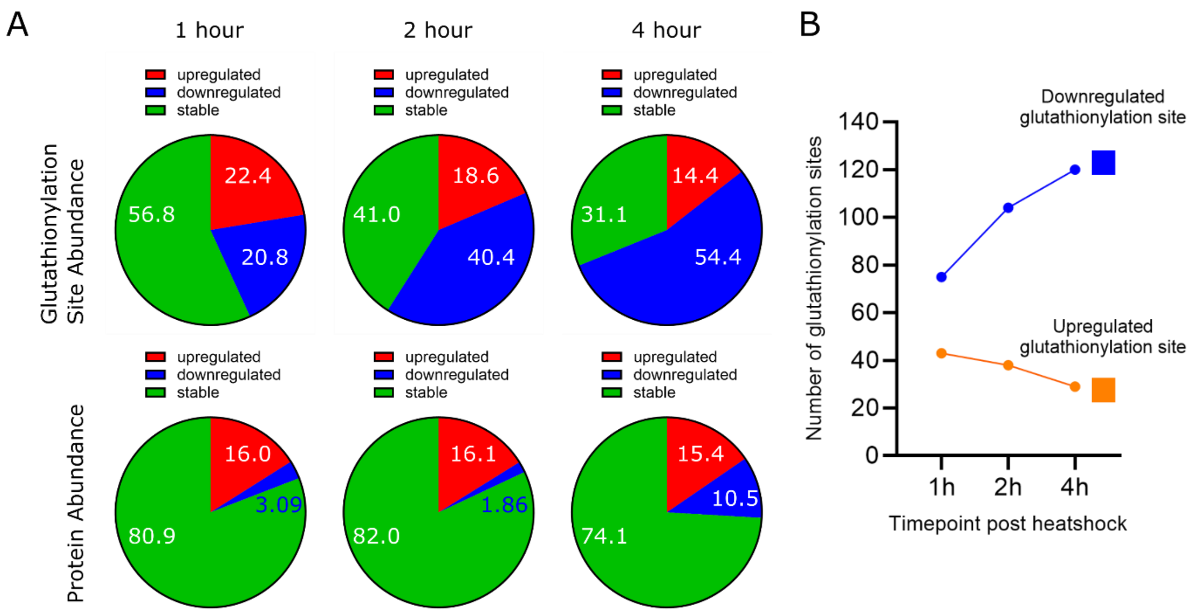


Figure S2. **Protein Glutathionylation is a dynamic PTM that responds to environmental stress**

(A) Venn diagrams to represent cysteine-glutathionylated peptide abundance ratio vs 0-hour control condition and glutathionylated protein abundance ratio vs 0-hour control condition. Upregulated glutathionylated peptide and proteins fulfilled a log2 fold change >1.5, downregulated glutathionylated peptide and proteins fulfilled a log2 fold change <-1.5, and stable proteins fulfilled a log2 fold change between -1.5 and 1.5 (all vs 0-hour condition). Numbers represent percentage of all glutathionylated peptides/proteins from that experimental condition.

(B) Number of protein-glutathionylations that are upregulated (orange) and downregulated (blue), at 1 h, 2 h and 4 h timepoints.

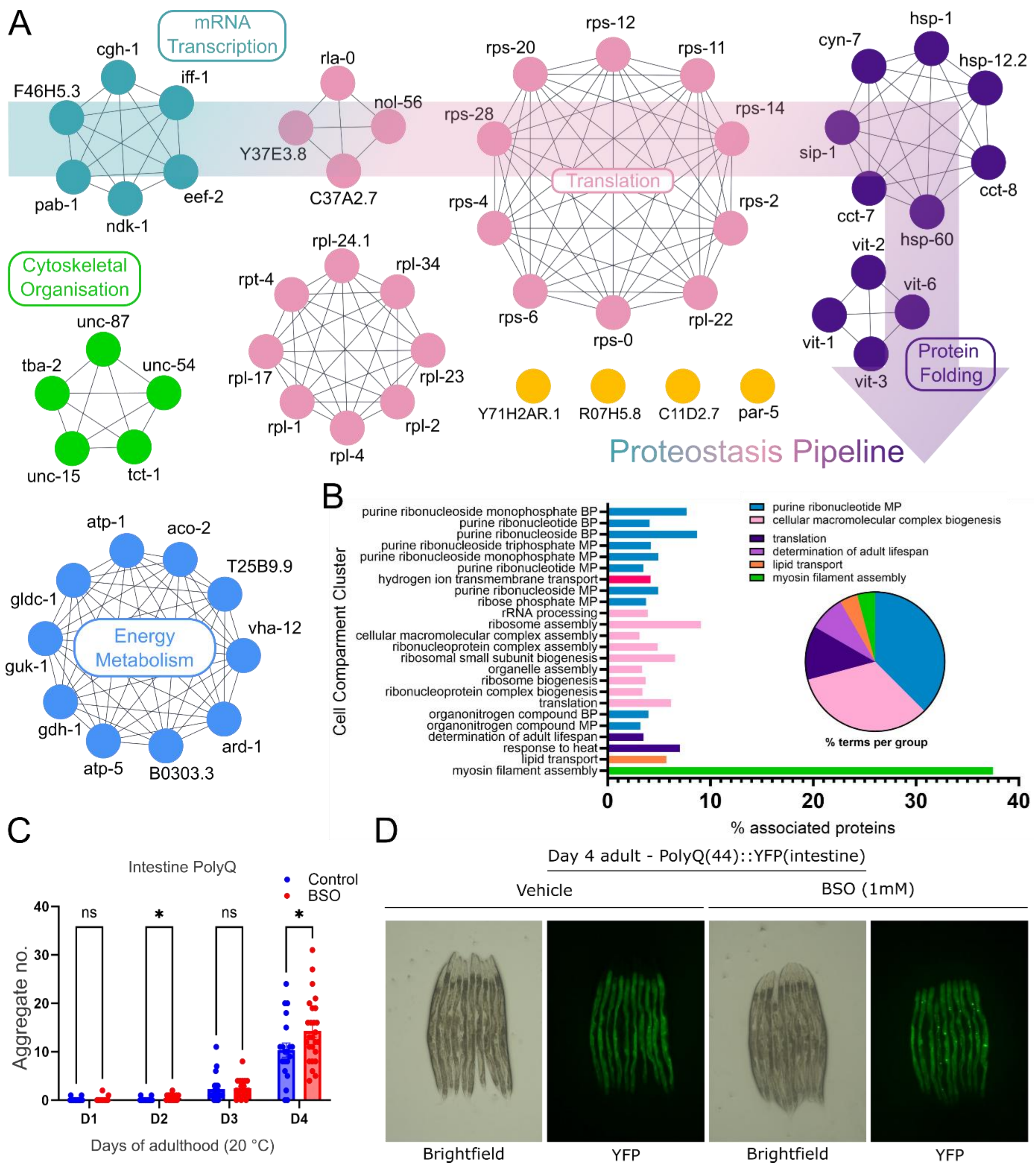


Figure S3: **Remodelling of protein-glutathionylation impacts protein homeostasis and tissue health.**

(A) Gene Ontology Biological Process analysis of proteins that underwent changes in glutathionylation upon genetic perturbation (Δ*daf2*) within *C. elegans*, relative to control.

(B) Gene Ontology Cell Compartment analysis of proteins that underwent changes in glutathionylation upon genetic perturbation (Δ*daf2*) within *C. elegans*, relative to control.

(C) Protein aggregation numbers for control vs BSO *C. elegans* intestine.

(D) Representative fluorescence microscopy images for intestine YFP-tagged polyQ *C. elegans*, control vs BSO.

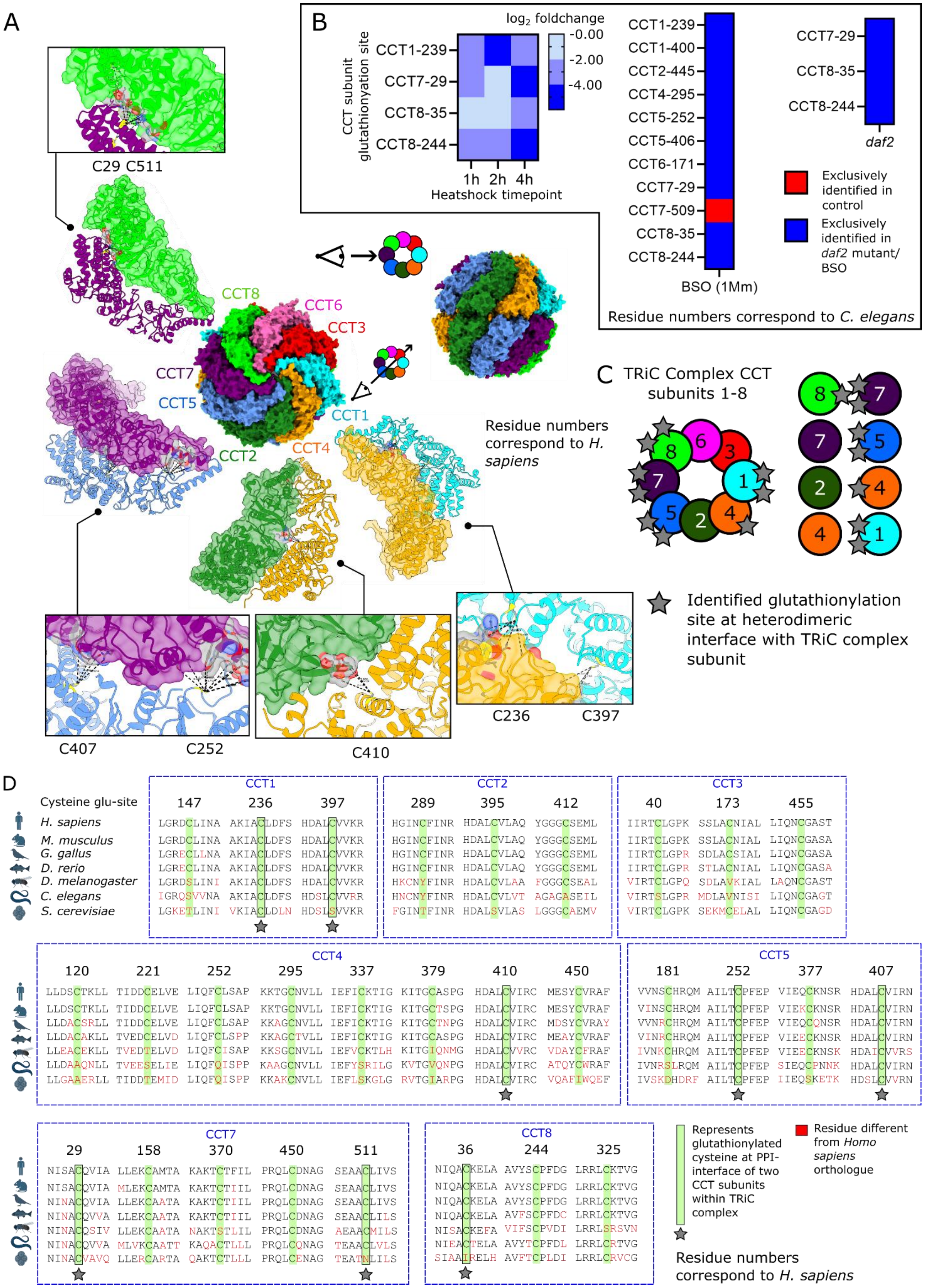

A
C29 C511
CCT8
CCT6
CCT3
CCT7
CCT5
CCT1
CCT2
CCT4
Residue numbers correspond to H. sapiens
C407 C252
C410
C236 C397
B
log2 foldchange
-0.00
-2.00
-4.00
CCT subunit glutathionyation site
CCT1-239
CCT7-29
CCT8-35
CCT8-244
1h 2h 4h
Heatshock timepoint
CCT1-239
CCT1-400
CCT2-445
CCT4-295
CCT5-252
CCT5-406
CCT6-171
CCT7-29
CCT7-509
CCT8-35
CCT8-244
BSO (1Mm)
CCT7-29
CCT8-35
CCT8-244
daf2
Exclusively identified in control
Exclusively identified in daf2 mutant/ BSO
Residue numbers correspond to C. elegans
C TRiC Complex CCT subunits 1-8
8 6 3 7 1 5 2 4
Identified glutathionylation site at heterodimeric interface with TRiC complex subunit
D
Cysteine glu-site
H. sapiens
M. musculus
G. gallus
D. rerio
D. melanogaster
C. elegans
S. cerevisiae
CCT1
147 236 397
LGRDCLINA AKIACLDFS HDALCVVKR
LGRDCLINA AKIACLDFS HDALCVVKR
LGRECLLNA AKIACLDFS HDALCVVKR
LGRECLINA AKIACLDFS HDALCVVKR
LGRDSLINI AKIACLDFS HDALCVVKR
IGRQSVVNA AKIACLDFS HDSLCVVRR
LGKETLINI VKIACLDLN HDSLSVVKR
CCT2
289 395 412
HGINCFINR HDALCVLAQ YGGGCSEML
HGINCFINR HDALCVLAQ YGGGCSEML
HGINCFINR HDALCVLAQ YGGGCSEML
HGINCFINR HDALCVLAQ YGGGCSEML
HKCNYFINR HDALCVLAA FGGGCSEAL
HNCNYFINR HDALCVLVT AGAGASEIL
FGINTFINR HDALSVLAS LGGGCAEMV
CCT3
40 173 455
IIRTCLGPK SSLACNIAL LIQNCGAST
IIRTCLGPK SSLACNIAL LIQNCGASA
IIRTCLGPR SDLACSIAL LIQNCGASA
IIRTCLGPR STLACNIAL LIQNCGASA
VIRTCLGPQ SDLAVKIAL LAQNCGAST
VIRTSLGPR MDLAVNISI LIQNCGAGT
VIRTCLGPK SEKMCELAL LIQNCGAGD
CCT4
120 221 252 295 337 379 410 450
LLDSCTKLL TIDDCELVE LIQFCLSAP KKTGCNVLL IEFICKTIG KITGCASPG HDALCVIRC MESYCVRAF
LLDSCTKLL TIDDCELVE LIQFCLSAP KKTGCNVLL IEFICKTIG KITGCTSPG HDALCVIRC MESYCVRAF
LLDACSRLL TIDDCELVE LIQFCLSAP KKAGCNVLL IEFICKTIG KITGCTNPG HDALCVIRC MDSYCVRAY
LLDACAKLL TVEDCELVD LIQFCLSPP KKAGCTVLL IEFICKTIG KITGCASPG HDALCVIRC MEAYCVRAF
LLEACEKLL TVEDTELVD LIQFCISAP KKSGCNVLL IEFVCKTLH KITGCTSPG HDALCVVRC VDAYCFRAF
LLDAAQNLL TVEESELIE LIQFQISPP KAAGCNVLL IEFYSRILG KVTGVQNPG HDALCVIRC ATQYCWRAF
LLGAAERLL TIDDTEMID LIQFQISPP KKAKCNVLL IEFLSKGLG RVTGIARPG HDALCVIRC VQAFIWQEF
CCT5
181 252 377 407
VVNSCHRQM AILTCPFEP VIEQCKNSR HDALCVIRN
VINSCHRQM AILTCPFEP VIEKCKNSR HDALCVIRN
VVNRCHRQM AILTCPFEP VIEQCQNSR HDALCVIRN
VINRCHRQM AILTCPFEP VIEECKNSR HDALCVIRN
IVNKCHRQM AILTCPFEP VIEECKNSK HDAICVVRS
IVNRSLRQM AILTCPFEP SIEQCPNNK HDALCVIRN
IVSKDHDRF AILTCPFEP IIEQSKETK HDSLCVVRN
CCT7
29 158 370 450 511
NISACQVIA LLEKCAMTA KAKTCTFIL PRQLCDNAG SEAACLIVS
NISACQVIA MLEKCAMTA KAKTCTIIL PRQLCDNAG SEAACLIVS
NINACQVIA LLEKCAATA KAKTCTIIL PRQLCDNAG SEAACLIVS
NINACQVVA LLEKCAATA KAKTCTIIL PRQLCDNAG SEAACLILS
NINACQSIV LLEKCAATA NAKTSTLIL PRQLCDNAG AEAACMILS
NINACQVVA MLVKCAATT KAQACTLLL PRQLCQNAG TEAACLVLS
NINACVAVQ LLERCARTA QAKTCTLLL PRQLCENAG TEATNLILS
CCT8
36 244 325
NIQACKELA AVYSCPFDG LRRLCKTVG
NIQACKELA AVYSCPFDG LRRLCKTVG
NIQACKELA AVYSCPFDG LRRLCKTVG
NIQACKELA AVFSCPFDC LRRLCKTVG
NISACKEFA VIFSCPVDI LRRLSRSVN
NIEACTELA AVYTCPFDL LRRLCRTVG
SIAAIRELH AVFTCPLDI LRRLCRVCG
Represents glutathionylated cysteine at PPI-interface of two CCT subunits within TRiC complex
Residue different from Homo sapiens orthologue
Residue numbers correspond to H. sapiens

Figure S4. **Remodelling of protein-glutathionylation impacts TRiC complex assembly**

(A) Crystal structure of the human TRiC complex (PDBe:7lup), with glutathionylation sites at protein-protein interfaces annotated with zoomed in structures, including: CCT1 cysteine-239 and -400 interfaced with CCT4, CCT4 cysteine-410 interfaced with CCT2, CCT5 cysteine-252 and -406 interfaced with CCT7, CCT7 cysteine-29 and -509 interfaced with CCT8 and CCT8 cysteine-35 interfaced with CCT7.

(B) Heatmaps to represent differential abundance of the 4, 11 and 3 heat-shock, BSO and *daf-2* dynamic glutathionylation sites.

(C) Schematic representation of glutathionylation across PPIs within the TRiC complex.

(D) Sequence conservation across Saccharomyces cerevisiae (yeast), *Caenorhabditis elegans* (worm), *Drosophila melanogaster* (fly), *Danio rerio* (fish), *Gallus gallus* (bird), *Mus Musculus* (rodent) and *Homo sapiens* (human) isoforms of each of the identified glutathionylated cysteines of CCT-1, -2, -3, -4, -5, -7 and -8. Experimentally identified glutathionylated cysteine site is annotated with green box and amino acids that are different vs *H. sapiens* orthologue are coloured red. Glutathionylation sites that are localised to protein-protein interfaces are annotated with a black outline around cysteine-site and star below box.

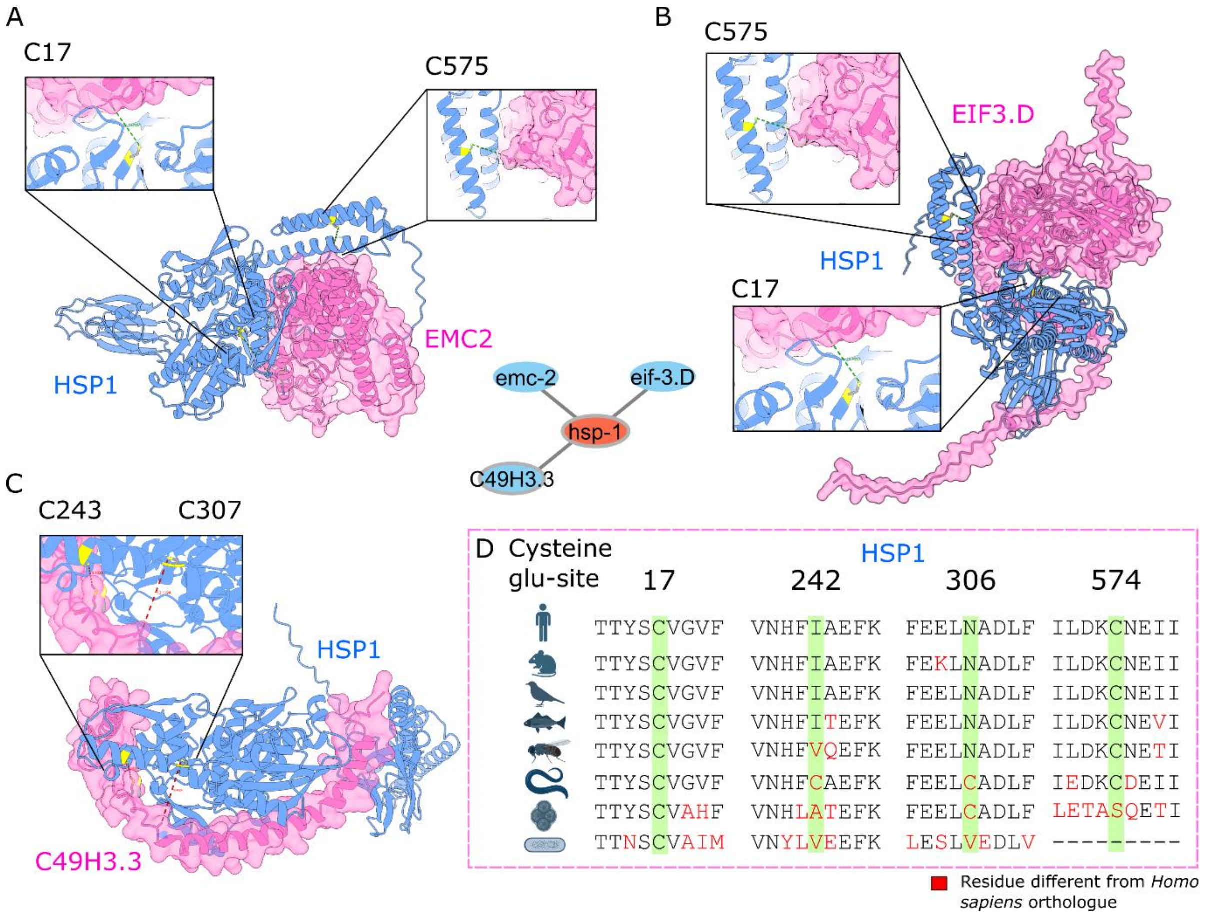


Figure S5: **Remodelling of protein-glutathionylation impacts HSP1 heterodimerization**

(A-C) AlphaFold generated heterodimeric PPIs of HSP1 (blue) with (A) EMC2, (B) EIF3.D and (C) C49H3.3, with annotated (yellow) dynamic glutathionylatable cysteine sites at the dimerization interface.

(D) Sequence conservation of HSP-1/HSC70 across *Escherichia coli* (DnaK), Saccharomyces cerevisiae (SSA3), *Caenorhabditis elegans (HSP-1)*, *Drosophila melanogaster* (*HSC70-4*), *Danio rerio (HSPA8)*, *Gallus gallus* (HSPA8), *Mus Musculus* (HSPA8) and *Homo sapiens (HSPA8)*. Experimentally identified glutathionylated cysteine site is annotated with green box and amino acids that are different vs *H. sapiens* orthologue are coloured red.

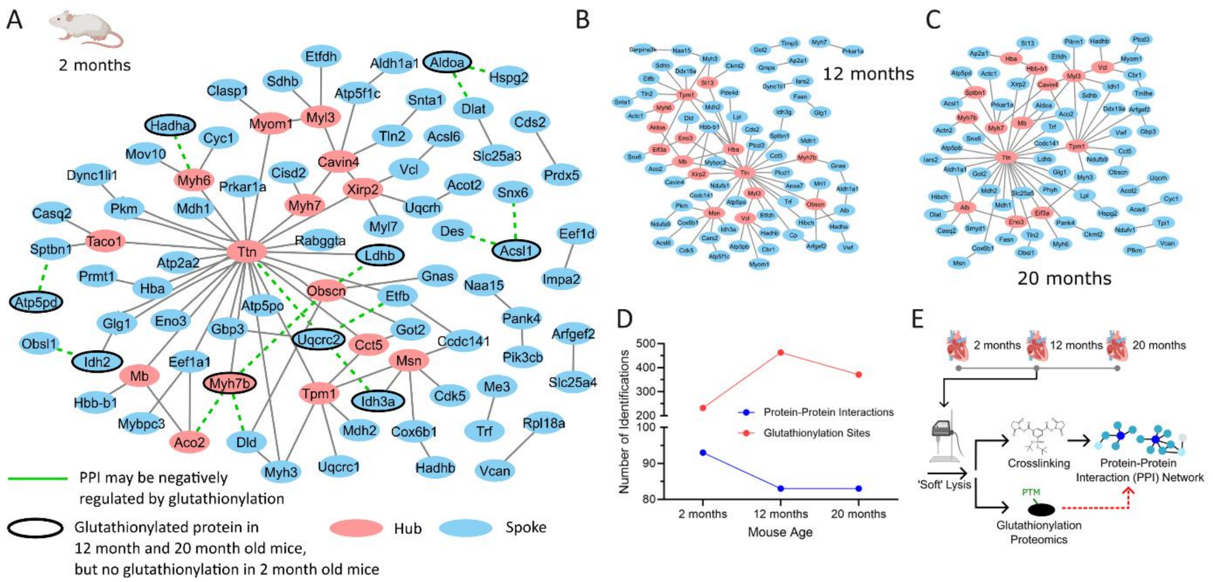


Figure S6. **Dynamic protein glutathionylation drives the redistribution of protein-protein interaction networks in mouse cardiac tissue**

(A-C) Protein-Protein interaction networks from whole cell crosslinking MS for (A) 2-month-old, (B) 12-month-old and (C) 20-month-old mice. For 2-month-old mice PPI network: proteins that were identified to be glutathionylated at 12- and 20-months old, but not at 2 months-old, are annotated with outline. PPI's that may be regulated by glutathionylation (as PPI was only observed when glutathionylation was absent) are annotated with green dashed lines.

(D) Graph to show the number of glutathionylation sites and number of protein-protein interactions for mouse cardiac tissue at three age groups.

(E) Workflow for glutathionylation and protein-protein interaction analysis of mouse cardiac tissue.

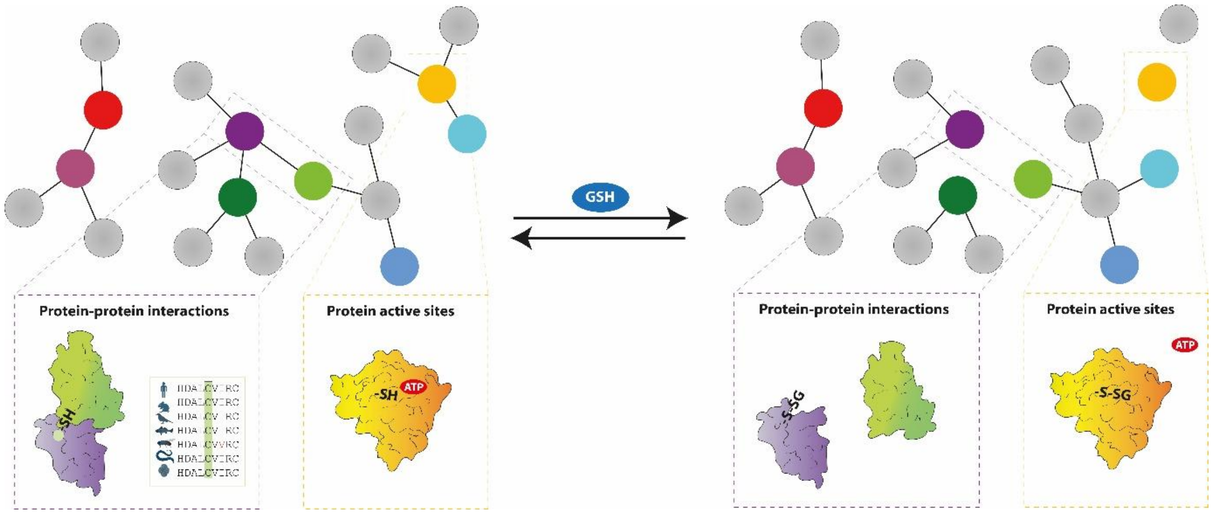


Figure S7. **Protein glutathionylation as a global regulator of protein-protein interactions and protein active site functionality.**
